# RNA velocity unraveled

**DOI:** 10.1101/2022.02.12.480214

**Authors:** Gennady Gorin, Meichen Fang, Tara Chari, Lior Pachter

**Affiliations:** Division of Chemistry and Chemical Engineering, California Institute of Technology, Pasadena, CA, 91125; Division of Biology and Biological Engineering, California Institute of Technology, Pasadena, CA, 91125; Department of Computing and Mathematical Sciences, California Institute of Technology, Pasadena, CA, 91125

## Abstract

We perform a thorough analysis of RNA velocity methods, with a view towards understanding the suitability of the various assumptions underlying popular implementations. In addition to providing a self-contained exposition of the underlying mathematics, we undertake simulations and perform controlled experiments on biological datasets to assess workflow sensitivity to parameter choices and underlying biology. Finally, we argue for a more rigorous approach to RNA velocity, and present a framework for Markovian analysis that points to directions for improvement and mitigation of current problems.

## 1 Introduction

### 1.1 Background

The method of *RNA velocity* [1] aims to infer directed differentiation trajectories from snapshot single-cell transcriptomic data. Although we cannot observe the transcription rate, we can count molecules of spliced and unspliced mRNA. The unspliced mRNA content is a leading indicator of spliced mRNA, meaning that it is a predictor of the spliced mRNA content in the cell’s near future. This causal relationship can be usefully exploited to identify directions of differentiation pathways without prior information about cell type relationships: “depletion” of nascent RNA suggests the gene is downregulated, whereas “accumulation” suggests it is upregulated. This qualitative premise has profound implications for the analysis of single-cell RNA sequencing (scRNA-seq) data. The experimentally observed transcriptome is a snapshot of a biological process. By carefully combining snapshot data with a causal model, it is for the first time possible to reconstruct the dynamics and direction of this process without prior knowledge or dedicated experiments.

The bioinformatics field has recognized this potential, widely adopting the method and generating numerous variations on the theme. The roots of the theoretical approach date to 2011 [2], but the two most popular implementations for scRNA-seq were released in 2017–2018: *velocyto* by La Manno et al. [1], which introduced the method, and *scVelo* by Bergen et al. [3], which extended it to fit a more sophisticated dynamical model. Aside from these packages, numerous auxiliary methods have been developed, including *protaccel* [4] for incorporating newly available protein data, *MultiVelo* [5] and *Chromatin Velocity* [6] for incorporating chromatin accessibility, *VeTra* [7], *CellPath* [8], *Cytopath* [9], *CellRank* [10], and *Revelio* [11] for investigating coarse-grained global trends, *scRegulosity* [12] for identifying local trends, *Dynamo* [13] for estimating the differentiation landscape curvature in metabolic labeling experiments, *Velo-Predictor* [14] for incorporating machine learning, *dyngen* [15] and *VeloSim* [16] for simulation, and *VeloViz* [17] and *evo-velocity* [18] for constructing velocity-inspired visualizations. This profusion of computational extensions has been accompanied by a much smaller volume of analytical work, including discussions of potential extensions and pitfalls [19–22], as well as theoretical studies based on optimal transport [23,24] and stochastic differential equations [25]. However, at their core, these auxiliary methods are built on top of the theory and code base from *velocyto* or *scVelo*.

These two most popular software implementations emphasize usability and integration with standard visualization methods. The typical user-facing workflows, with internal logic abstracted away, are shown in Figure 1: a set of reads is converted to cell × gene matrices derived from spliced and unspliced mRNA molecule measurements, the matrices are processed to generate phase plots describing a dynamical transcription process, and finally the transcriptional dynamics are fit, extrapolated, and displayed in a low-dimensional embedding.

**Figure 1:**
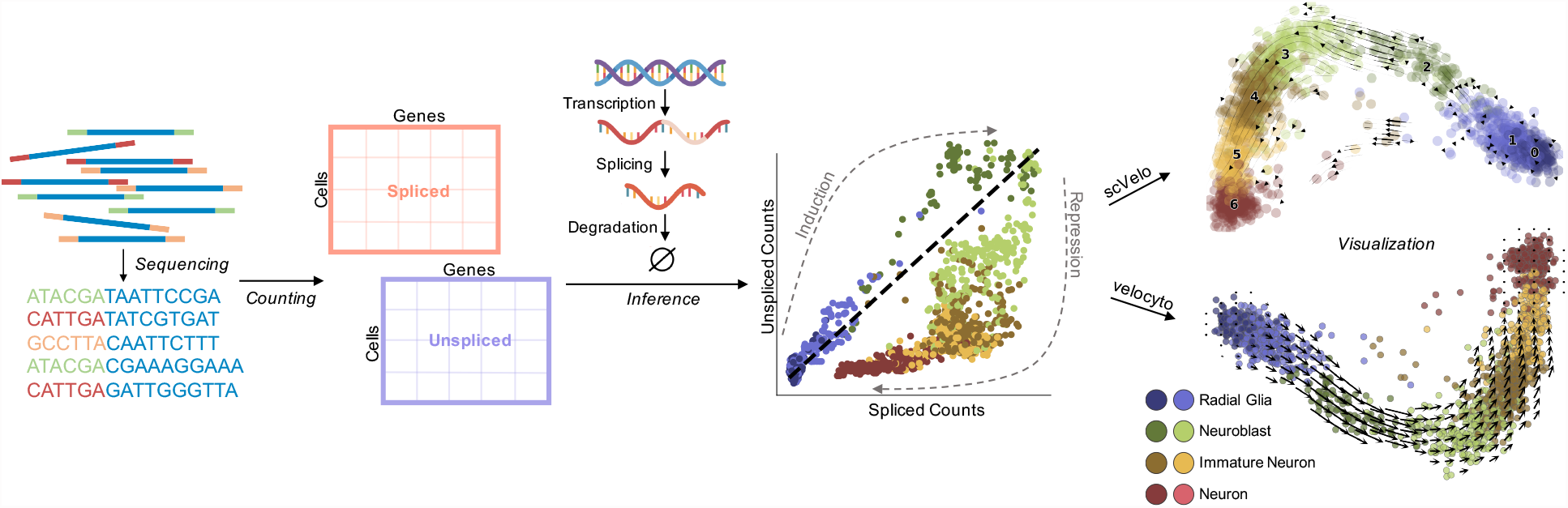
A summary of the user-facing workflow of a typical RNA velocity workflow. Initial processing of sequencing reads produces spliced and unspliced counts for every cell, across all genes. Inference procedures, implemented in *velocyto* and *scVelo*, fit a model of transcription, and predict cell-level velocities. The final embedding of cells and smoothed velocities is displayed in the top two principal component dimensions. Visualizations adapted from [26, 27]; dataset from [1].

Despite the popularity of RNA velocity [14, 28] and increasingly sophisticated attempts to combine it with more traditional methods for trajectory inference [8, 10], there has been no comprehensive investigation of the modeling assumptions that underlie the seemingly simple workflow. This is an impediment to applying, interpreting, and refining the methods, as problems arise even in the simplest cases. Consider, for example, the result displayed in Figure 1, where the outputs of the two most popular RNA velocity programs applied to human embryonic forebrain data generated by La Manno et al. [1] (“forebrain data”) are qualitatively different. The inferred directions in the example should recapitulate a known differentiation trajectory from radial glia to mature neurons. However, *scVelo*, which “generalizes” *velocyto*, fails to identify, and even reverses the trajectory, suggesting totally different causal relationships between cell types. This type of problematic result has been reported elsewhere (Figs. 2-3 of [3], Fig. 2 of [22], Fig. 4B of [5], Fig. 5A of [9], Fig. 5 of [10], and Fig. 3 of [13]).

**Figure 2:**
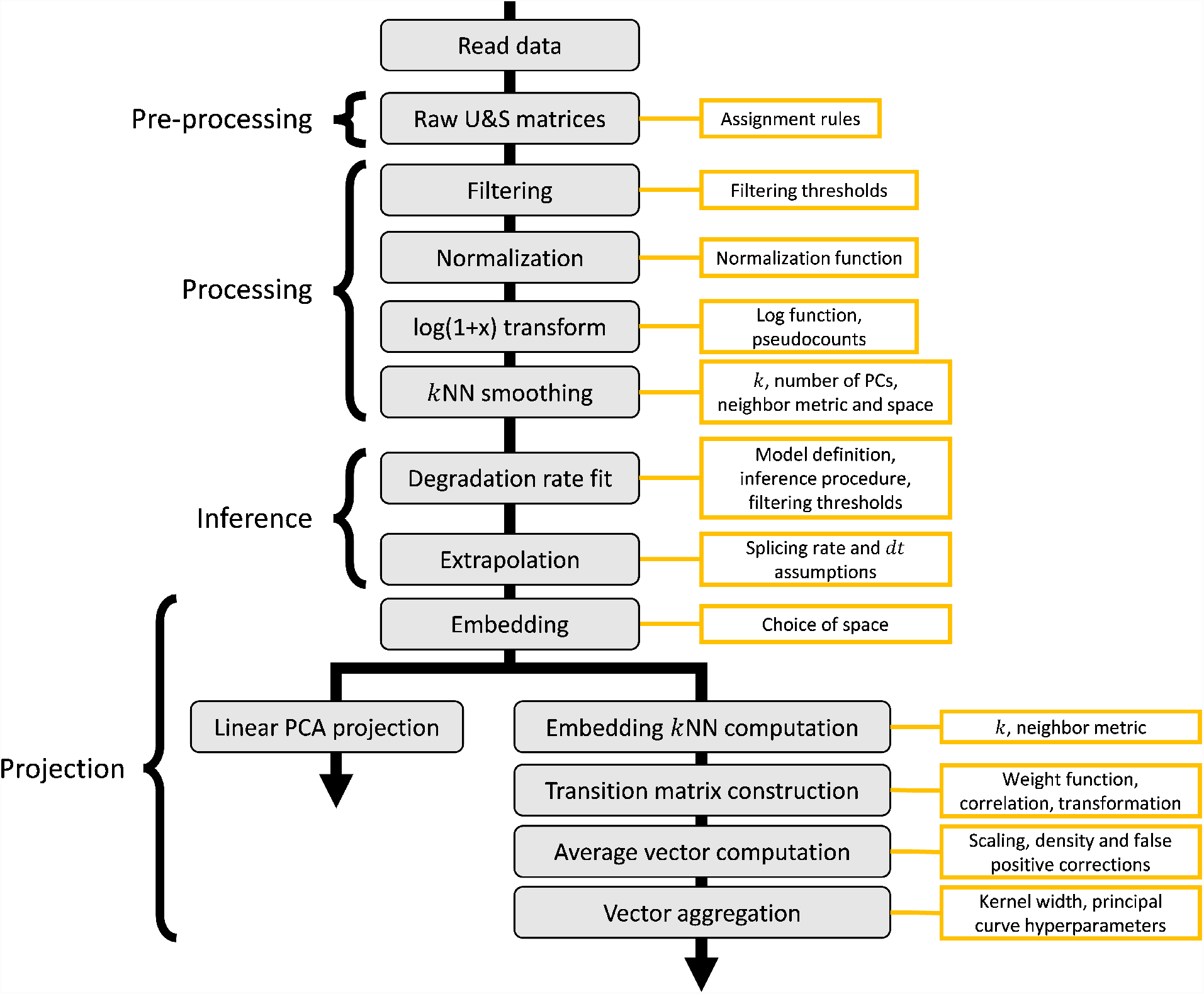
An RNA velocity workflow, beginning with read processing and ending with twodimensional projection, and the parameters that must be specified by the user.

**Figure 3:**
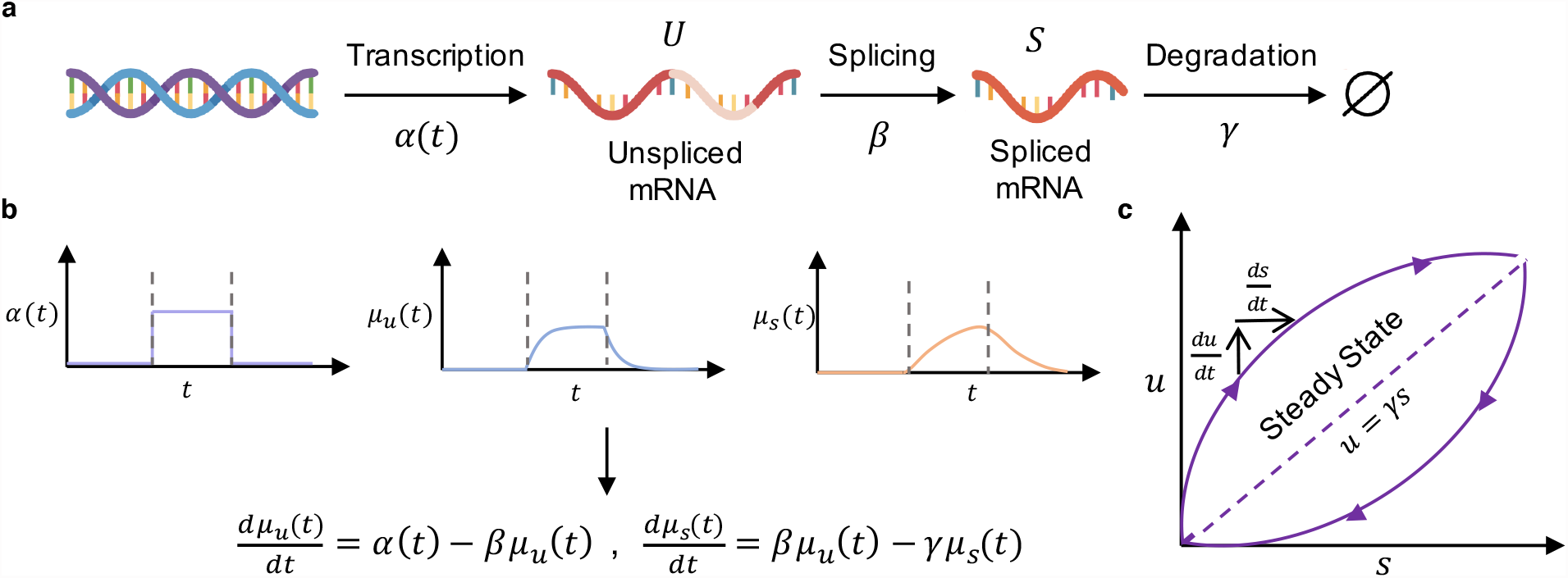
**a**. The continuous model of transcription, splicing, and degradation used for RNA velocity analysis. **b**. Plots of *α*(*t*), *μ*_*u*_(*t*), and *μ*_*s*_(*t*) over time *t* and the corresponding governing equations for the system. Dashed lines indicate time of transcription event. **c**. Outline of the common phase portrait representation, with both steady state and dynamical models denoted. Adapted from [1]

**Figure 4:**
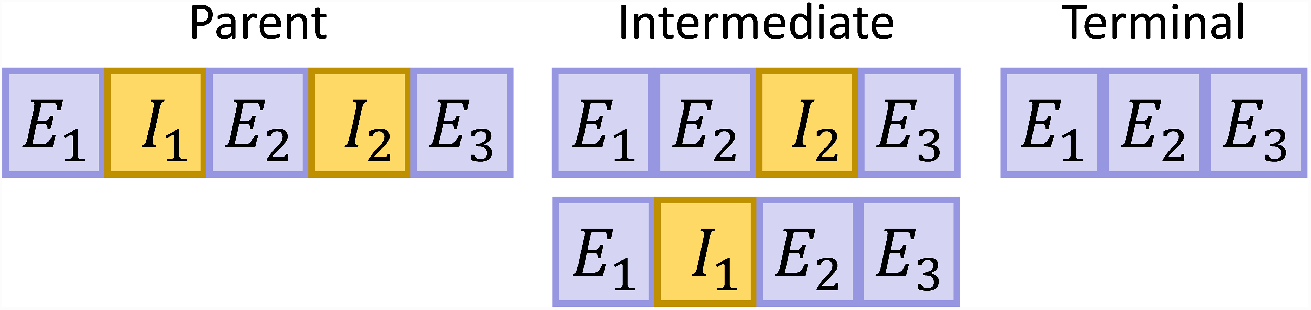
A two-intron mRNA species may not have well-defined “unspliced” and “spliced” forms.

**Figure 5:**
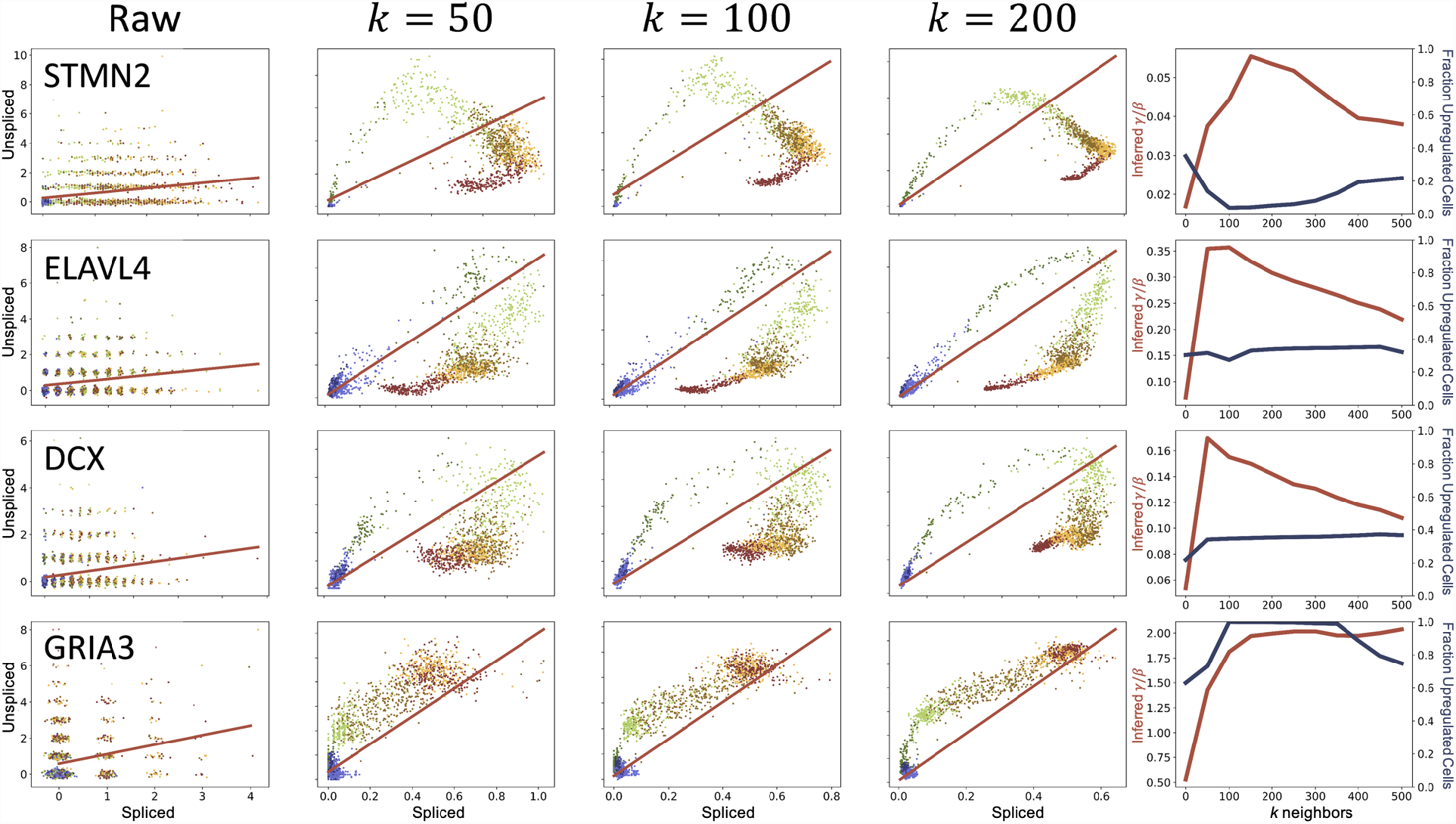
Distortions in data and instabilities in the inferred *γ/β* values introduced by the imputation procedure on the forebrain data from [1]. Column 1: Raw data (points: spliced and unspliced counts with added jitter; color: cell type, as in Figure 1; line: best fit line *u* = *γs/β* + *q*, estimated from the entire dataset). Columns 2-4: Normalized and imputed data under various values of *k* (points: spliced and unspliced counts; color: cell type, as in Figure 1; line: best linear fit *u* = *γs/β* + *q*, estimated from extreme quantiles). Column 5: Inferred values of *γ/β* (red, left axis) and inferred fraction of upregulated cells, defined as Σ_*i*_ *u*_*i*_ − (*γs*_*i*_*/β* + *q*) > 0 (blue, right axis).

Motivated by such discrepancies, we wondered whether either *velocyto* or *scVelo* are reliable for standard use in applications where ground truth may be unknown. An examination of their theoretical foundations, and those of related methods, revealed that they are largely informal. Even the term “RNA velocity” is not precisely defined, and is used for the following distinct concepts:

- A generic method to infer trajectories and their direction using relative unspliced and spliced mRNA abundances by leveraging the causal relationship between the two RNA species, which is the interpretation in [25].
- A set of tools implementing this method or parts of it, as in an “RNA velocity workflow implemented in *kallisto*|*bustools*”, which is the interpretation in [29].
- A gene- and cell-specific quantity under a continuous model of transcription, as in “the RNA velocity of a cell is 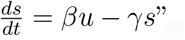, which is the interpretation in [19, 28].
- A gene- and cell-specific quantity under a probabilistic model of transcription, as in “the RNA velocity of a cell is 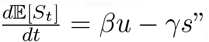, which is the interpretation in [4].
- A gene-specific average quantity, as in “the total RNA velocity of a gene is Σ_*i*_(*βu*_*i*_ − *γs*_*i*_)”, which is the interpretation in [12, 28].
- A cell-specific vector composed of gene-specific velocity components, as in “the vector RNA velocity of a cell is *β*_*j*_*u*_*ij*_ − *γ*_*j*_*s*_*ij*_”, which is the interpretation in [7, 9, 28].
- The cell-specific linear or nonlinear embedding of a cell-specific vector in a low-dimensional space, which is the interpretation in [9].
- A local property, such as curvature, of a theorized cell landscape computed either from an embedding or a set of velocities, which is the interpretation in [13, 23].

These discrepancies and, more broadly, the limitations of current theory, stem from historical differences between sub-fields, which have calcified over the past twenty years of single-cell biology. On the one hand, fluorescence transcriptomics methods, including single-molecule fluorescence *in situ* hybridization and live-cell MS2 tagging, which target small, well-defined systems with a narrow set of probes [30–32], have motivated the development of interpretable stochastic models of biological variation [33, 34]. On the other hand, “sequence census” methods [35], such as scRNA-seq, provide genome-wide quantification of RNA, but the associated challenges of exploratory, high-dimensional data analysis have not, for the most part, been addressed with mechanistic models. Instead, descriptive summaries, such as graph representations and low-dimensional embeddings, are the methods of choice [36]. Nevertheless, descriptive analyses, even if *ad hoc*, can still facilitate biological discovery: RNA velocity has been used to produce plausible trajectories [37–45], and our simulations show that it can recapitulate key information about differentiation trajectories in best-case scenarios (Figure S1). These results highlight the potential of RNA velocity, and motivated us to review its assumptions, understand its current failure modes, and to solidify its foundations.

Towards this end, we found it helpful to contrast the sub-fields of fluorescent transcriptomics and sequencing, which have analogous goals, albeit disparate origins that have led to analytical methods with distinct philosophies and mathematical foundations. The sub-fields have, at times, interacted. Fluorescence transcriptomics can now quantify thousands of genes at a time, and this scale of data is now occasionally presented using visual summaries popular for RNA sequencing data, such as principal component analysis (PCA) [46], Uniform Manifold Approximation and Projection (UMAP) [47], and *t*-distributed stochastic neighbor embedding (*t*-SNE) [48,49]. Conversely, the commercial introduction of scRNA-seq protocols with unique molecular identifiers (UMIs) has spurred the adoption of theoretical results from fluorescence transcriptomics for sequence census analysis [50–54]. Sequencing studies frequently use count distribution models that arise from stochastic processes, such as the negative binomial distribution, albeit without explicit derivations or claims about the data-generating mechanism [50,55,56]. These connections highlight the promise of mechanistic gene expression models: in principle, parameters can be fit to sequencing data to produce a physically interpretable, genome-scale model of transcriptional regulation in living cells, and some steps have been taken in this direction over the past decade [51–54, 57].

RNA velocity methods are products of the sequence census paradigm: they draw heavily on low-dimensional embeddings and graphs derived from the raw data. Their current limitations stem from viewing biology through the lens of signal processing, where noise is something to be eliminated or smoothed out. We posit that it is more appropriate to view the data through the lens of quantitative fluorescence transcriptomics, in which noise is a biophysical phenomenon in its own right. Through this lens, modeling that decomposes variation into single-molecule (intrinsic) and cell-to-cell (extrinsic) [58] components, in addition to technical noise [59], is key. Beyond this conceptual issue, we find that an assessment of the impact of hyper-parameterized, heuristic data pre-processing and visualization in current RNA velocity workflows is useful for developing more reliable analyses.

### 1.2 Goals and findings

To fully describe what RNA velocity does, why it may fail, and how it can be improved, requires work on several fronts:

In Section 2, we describe an idealized “standard” RNA velocity workflow. We introduce the biophysical foundations presented in the original publication, outline the methodological choices implemented in the software packages, and enumerate the tunable hyperparameters left to the user.

In Section 3, we probe the logic of the assumptions made in the workflow and describe potential failure points. This analysis revisits the outline through complementary critical lenses, adapted to the mechanistic and phenomenological steps. To characterize its biological coherence, we compare the concrete and implicit biophysical models to those standard in the field of fluorescence transcriptomics, and discuss the implications of assumptions that do not appear to be backed by a biophysical or mathematical argument. To characterize its stability, we test the quantitative effects of tuning hyperparameters and using different software implementations on real datasets.

Our findings on RNA velocity have implications for other scRNA-seq analyses. On one hand, the theory behind RNA velocity is not sufficiently robust. The models disagree with known biophysics: they do not recapitulate bursty production, and place needlessly restrictive constraints on regulatory trends. They are also internally inconsistent, as they do not preserve cell identities. Furthermore, the embedding processes are *ad hoc* and heavily rely on error cancellation, apparently discarding much of the data in the process. These problems are intrinsic, and derived methods inherit them.

Fortunately, better options, inspired by fluorescence transcriptomics models, are available. In order to develop a meaningful foundation for RNA velocity, we formalize its stochastic model and describe an inferential procedure that can be internally coherent and consistent with transcriptional biophysics. Furthermore, by examining the assumptions underpinning RNA velocity and reframing them in terms of stochastic dynamics, we find that the *velocyto* and *scVelo* procedures naturally emerge as approximations to our solutions. Our approach, presented in Section 4, provides an alternative to current trajectory inference methods: instead of using physically uninterpretable adjacency metrics and fitting a narrow set of topologies, it is relatively straightforward to solve any combination of transient or stationary topologies and apply standard Bayesian methods to identify the best fit. Conceptually, instead of “denoising” data, our approach proposes fitting the molecule distributions and preserving the uncertainty inherent in noisy biological and experimental processes.

## 2 Workflow and implementations

We begin with a conceptual overview of an idealized RNA velocity workflow, with a description of implementation-specific choices. We focus on datasets with cell barcodes and UMIs, such as those generated by the 10x Genomics Chromium platform [60], as they provide the most natural comparison to discrete stochastic models later in the discussion (Section 4.2). We summarize the workflow in Fig. 2, giving particular attention to the parameter choices required at each step. To clarify the information transfer in the process, we report the manipulations performed and the variables defined in a single run of the processing workflow in Fig. S2 (as used to generate Fig. 4 of [1]).

### 2.1 Pre-processing

RNA velocity analysis begins by processing raw sequencing data to distinguish spliced and unspliced molecules. This is a genomic alignment problem. For example, reads aligning to intronic references are assigned to unspliced molecules, whereas reads spanning exon-exon splice junctions are assigned to spliced molecules. Data from reads associated with a single UMI are combined to generate an label of “spliced,” “unspliced,” or “ambiguous” for each read. “Ambiguous” reads are omitted from downstream analysis, so the assignments are effectively binary.

Until recently, traditional alignment and UMI counting software, such as *Cell Ranger* from 10x Genomics, discarded intronic information [60]. The same was true of pseudoalignment methods, as they identify transcript classes consisting of annotated, and presumably terminal, isoforms [61]. The explicit quantification of transient intron-containing molecules appears to have been introduced in the *velocyto* command-line interface [1]. Since then, existing workflows have added functionality for unspliced transcript quantification [28]. In particular, alignment can be performed via *STAR-solo* [62] and *dropEst* [63], whereas pseudoalignment can be performed via *kallisto*|*bustools* [29] or *salmon* [28]. Benchmarking has shown discrepancies between the outputs of these workflows [28,29], apparently due to differences in filtering, thresholding, and counting ambiguous reads. However, there is currently little principled reason to prefer one program’s results to another, as quantification rules largely follow *velocyto*, and assume a two-species model is sufficient.

### 2.2 Count processing

The raw count data are processed to smooth out noise contributions that can skew the downstream analysis. This step is generally combined with the standard quality control techniques for scRNA-seq [36]. First, cells with extremely low expression are filtered out. Then, a subset of several thousand genes with the highest expression and variation are selected. The counts are normalized by the number of cell UMIs to counteract technical and cell size effects. At this point, the PCA projection is computed from log-transformed spliced RNA counts. Finally, the normalized counts are smoothed out by nearest-neighbor pooling. To accomplish this, the algorithm computes the *k* nearest cell neighbors in a PCA space for each cell, then replaces the abundance with the neighbors’ average. This step is crucial, as it produces the cyclic or near-cyclic “phase portraits” used in the inference procedure.

The implementation specifics vary even between the two most popular packages, the Python versions of *velocyto* and *scVelo*. For example, there appears to be no consensus on the appropriate *k* or neighborhood definition for imputation. The original publication reports *k* between 5 and 550, calculated using Euclidean distance in 5-19 top PC dimensions [1]. By default, *scVelo* uses *k* = 30 in the top 30 PC dimensions [3].

### 2.3 Inference

The normalized and smoothed count matrices are fit to a biophysical model of transcription. The model structure for a single gene is outlined in Figure 3a. *α*(*t*) is a transcription rate, which has pulse-like behavior over the course of the trajectory. The constant parameters are *β*, the splicing rate, and *γ*, the degradation rate. Driving by *α*(*t*) induces continuous trajectories *μ*_*u*_(*t*) and *μ*_*s*_(*t*), which informally represent instantaneous averages, *μ*, of the unspliced, *u*, and spliced, *s*, species, governed by the following ordinary differential equations (ODEs):

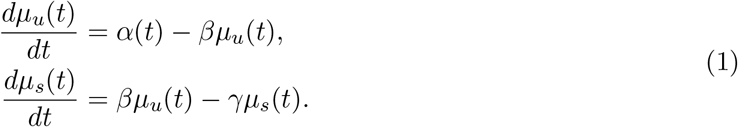

The qualitative behaviors of these functions are shown in Figure 3b. By fitting smoothed count data for a single gene, now interpreted as samples from a dynamical phase portrait governed by Equation 1 (Figure 3c), it is possible to estimate the ratio *γ/β*. Finally, with this ratio in hand, the velocity *v*_*i*_ may be computed for each cell *i*:

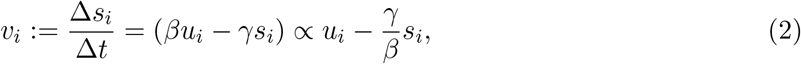

where *s*_*i*_ and *u*_*i*_ are cell-specific counts, Δ*t* is an arbitrary small time increment, and Δ*s*_*i*_ is the change in spliced counts achieved over that increment.

The popular packages differ on the appropriate way to fit the rate parameters. The *velocyto* procedure presupposes that the system reaches equilibria at the low- and high-expression states of *α*(*t*), and approximates them by the extreme quantiles of the phase plots. By computing the slope of a linear fit to these quantiles, it obtains the parameter *γ/β* (Fig. 3c). On the other hand, *scVelo* relaxes the assumption of equilibrium and implements a “dynamical” model, which fits the solution of Equation 1 to the entire phase portrait to obtain *γ* and *β* separately. This methodological difference corresponds to conceptual differences in the interpretation of imputed data. In *velocyto*, imputation appears to be an *ad hoc* procedure for filtering technical effects, in line with the usual usage [64, 65]. On the other hand, in *scVelo*, the imputed data are called “moments” and treated as identical to the instantaneous averages *μ*_*u*_(*t*) and *μ*_*s*_(*t*) of the process. In addition, *scVelo* offers a “stochastic” model, which posits pooled second moments are equivalent to the instantaneous second moments (e.g., the sum of *s*^2^ over neighbors is equal to 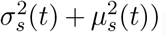).

The genes are analyzed independently, generating a velocity *v*_*ij*_ for each cell *i* and gene *j*. As the *velocyto* procedure cannot separately fit *β*_*j*_ and *γ*_*j*_, its velocities have different units for different genes. On the other hand, the *scVelo* procedure does separately fit the rate parameters, albeit by assigning a latent time *t*_*ij*_ to each cell, distinct for each gene’s fit.

### 2.4 Embedding

Low-dimensional representations are generated using one of the conventional algorithms, such as PCA, *t*-SNE, or UMAP. These algorithms can be conceptualized as functions that map from a high-dimensional vector *s*_*i*_ to a low-dimensional vector *E*(*s*_*i*_). The original publication offers two methods to convert cell’s velocity vector *v*_*i*_ to a low-dimensional representation [1].

If the embedding is deterministic (e.g., *E* is PCA on log-transformed counts), one can define a source point *E*(*s*_*i*_), compute a destination point *E*(*s*_*i*_ + *v*_*i*_Δ*t*) = *E*(*s*_*i*_ + Δ*s*_*i*_), and take the difference of these two low-dimensional vectors to obtain a local vector displacement:

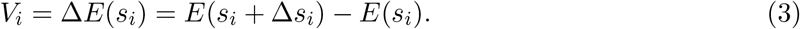

This displacement is then interpreted as a scalar multiple of the cell-specific embedded velocity.

If the embedding is non-deterministic, one can apply an *ad hoc* nonlinear procedure. This procedure essentially computes an expected embedded vector by weighting the directions to *k* embedding neighbors; neighbors that align with Δ*s*_*i*_ are considered likely destinations for cell state transitions in the near future:

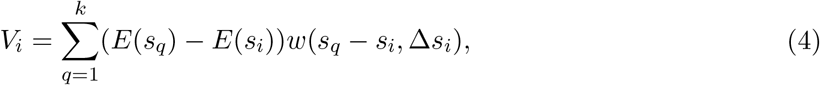

where *w* is a composition of the softmax operator (with a tunable kernel width parameter) with a measure of concordance between the arguments. Once an average direction is computed, it undergoes a set of corrections, e.g., to remove bias toward high-density regions in the embedded space. Finally, the cell-specific embedded vectors are aggregated to find the average direction over a region of the low-dimensional projection.

The packages almost exclusively use the nonlinear embedding procedure. There is no consensus on the appropriate choice of embedding, number of neighbors, or measure of concordance. PCA, *t*-SNE, and UMAP have been used to generate low-dimensional visualizations [1, 3]. The original publication uses *k* between 5 and 300 and applies square-root or logarithmic transformations prior to computing the Pearson correlation between the velocity and neighbor directions [1]. In contrast, *scVelo* uses a recursive neighbor search by averaging over neighbors and neighbors of neighbors (with *k* = 30), and implements several variants of cosine similarity [3]. An optional step adjusts the embedded velocities by subtracting a randomized control; this correction is usually omitted in demonstrations of *velocyto* and implemented but apparently undocumented in *scVelo*.

As demonstrated in Figure 2, the linear PCA embedding is the simplest dimensionality reduction technique; it consists of a projection and requires fewer parameter choices than other methods. However, it is only consistently used in *Revelio* [11]. The *velocyto* package does not appear to have a native implementation of this procedure, although it is briefly demonstrated in the original article (Fig. 2d and SN2 Figs. 8–9 of [1]). On the other hand, *scVelo* does implement the PCA velocity projection, but disclaims the results of using it as unrepresentative of the high-dimensional dynamics.

**Figure 6:**
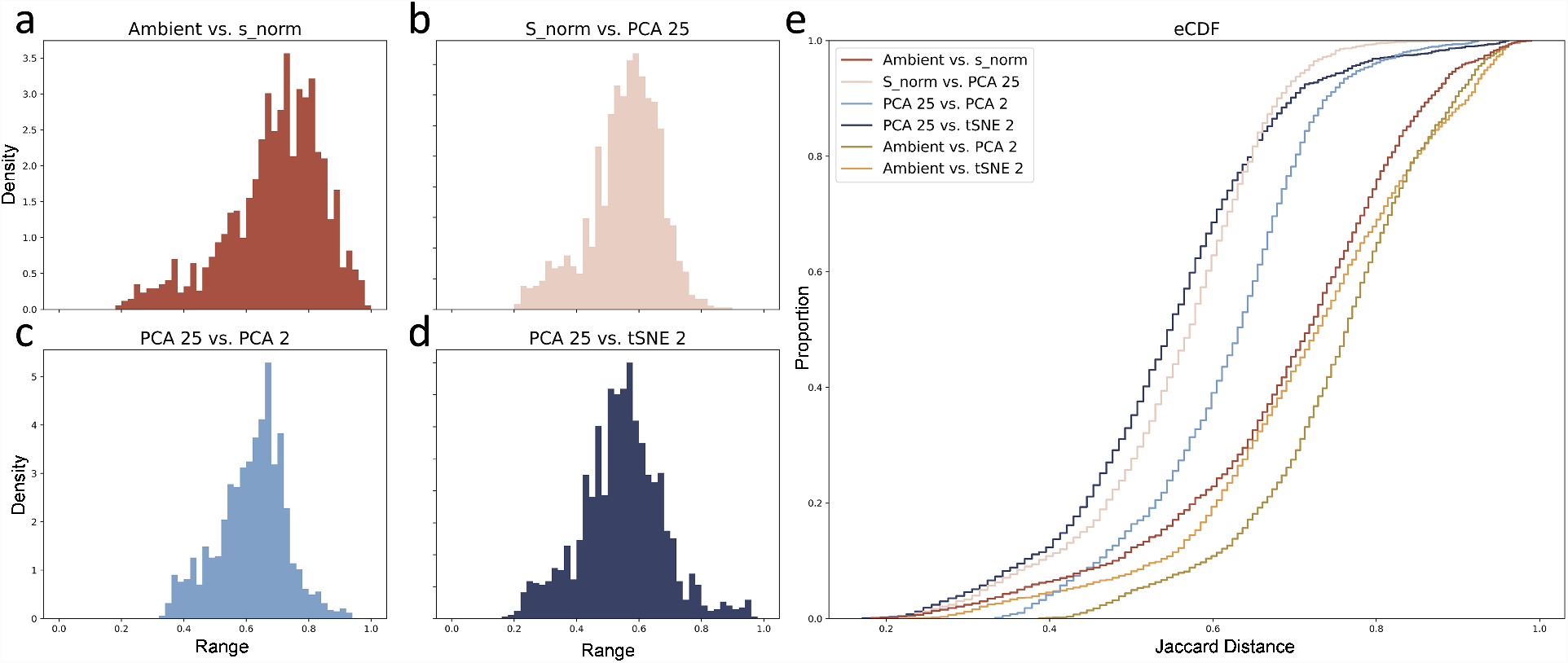
Normalization followed by two rounds of dimensionality reduction introduce distortions in the local neighborhoods. **a**.–**d**. Histograms of Jaccard distances between intermediate embeddings. **e**. Empirical cumulative distribution functions of Jaccard distances between intermediate embeddings, as well as the overall distortion (Ambient vs. PCA 2 and Ambient vs. tSNE 2).

**Figure 7:**
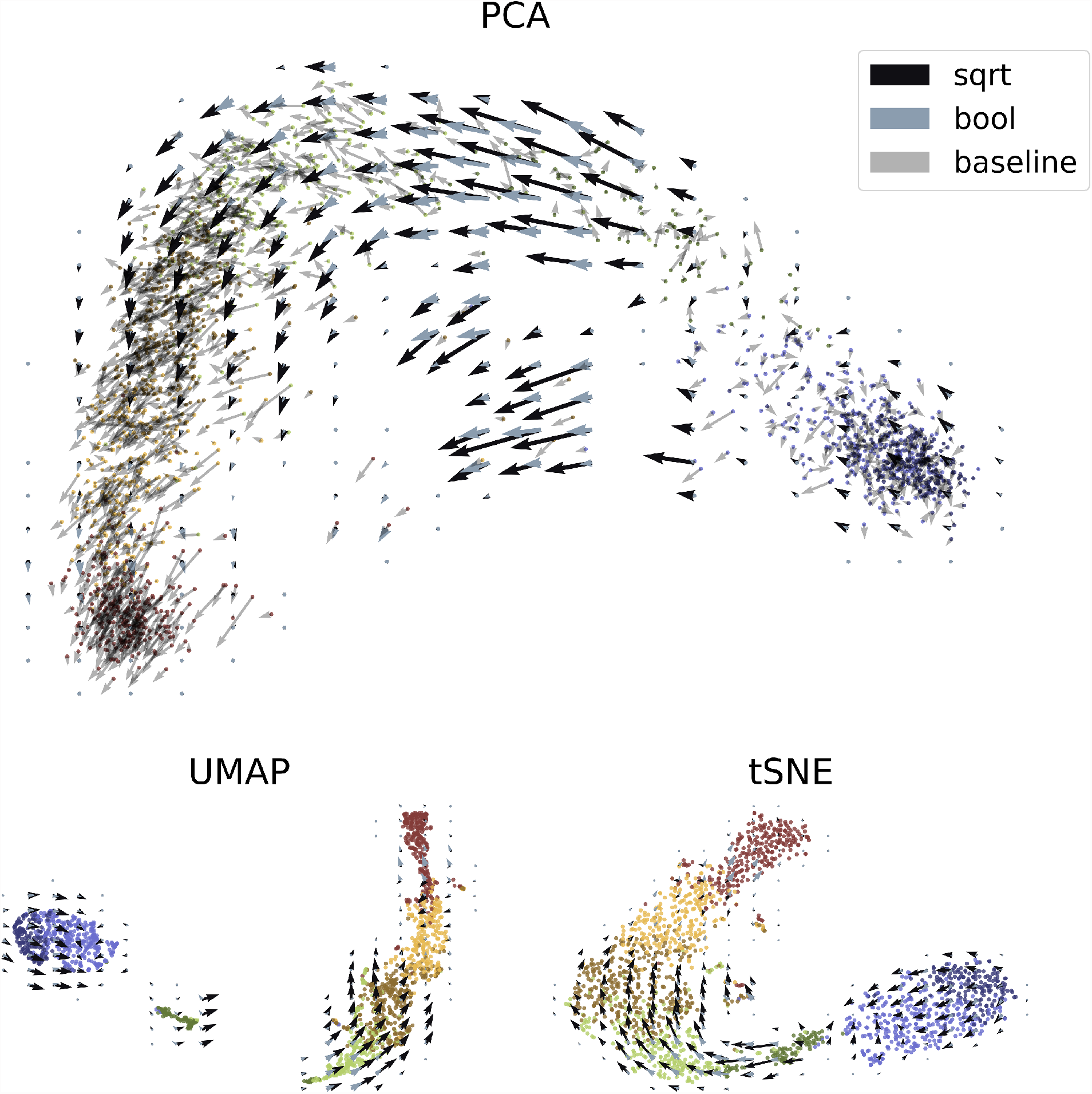
Performance of cell and velocity embeddings on the forebrain data. Top: PCA embedding with linear baseline and nonlinear aggregated velocity directions. Bottom: UMAP and *t*-SNE embeddings with nonlinear velocity projections.

**Figure 8:**
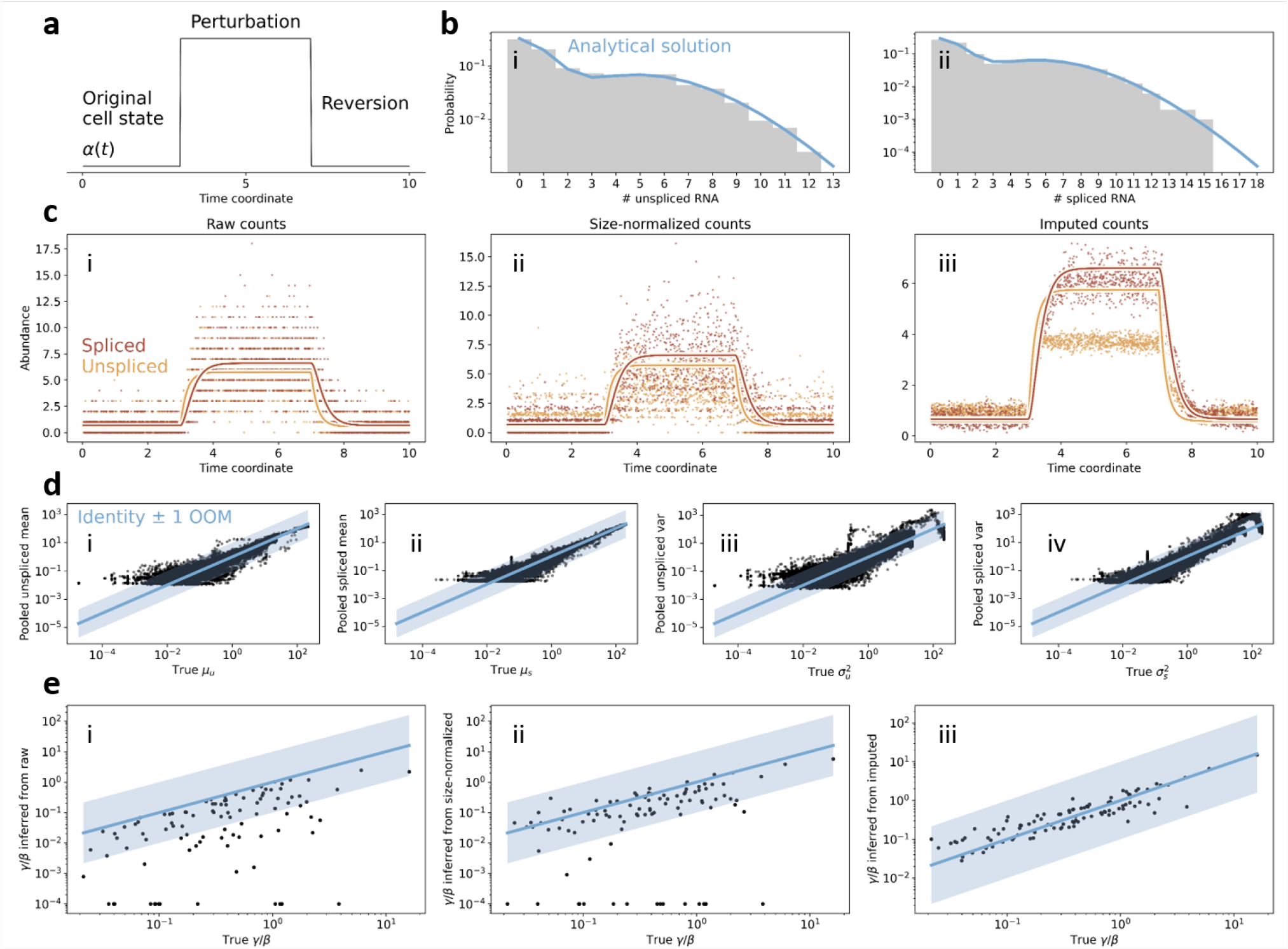
The RNA velocity count processing and inference workflow, applied to data generated by stochastic simulation. **a**. Schematic of the impulse model of gene modulation. **b**. Demonstration of the concordance between simulation and analytical solution for the occupation measure. **i**.: nascent mRNA counts; **ii**.: mature mRNA counts (gray: simulation; blue: occupation measure). **b**. Smoothing and imputation introduce distortions into the data. **i**.: raw data; **ii**.: data normalized to total counts; **iii**. imputed data (points: raw or processed observations; lines: ground truth averages *μ*_*s*_ and *μ*_*u*_; red: spliced; yellow: unspliced). **c**. Local averages obtained by imputation are not interpretable as instantaneous averages. **i**.: mean unspliced; **ii**.: mean spliced; **iii**. variance unspliced; **iv**.: variance unspliced (black points: true moment vs. pooled moment; blue line: identity; blue region: factors of ten around identity). **d**. Smoothing and imputation improve the inference on extrema. **i**.: moment-based inference from raw data; **ii**.: extremal inference from normalized data; **iii**.: extremal inference from imputed data (black points: true vs. inferred values of *γ/β*; blue line: identity; blue region: factors of ten around identity).

**Figure 9:**
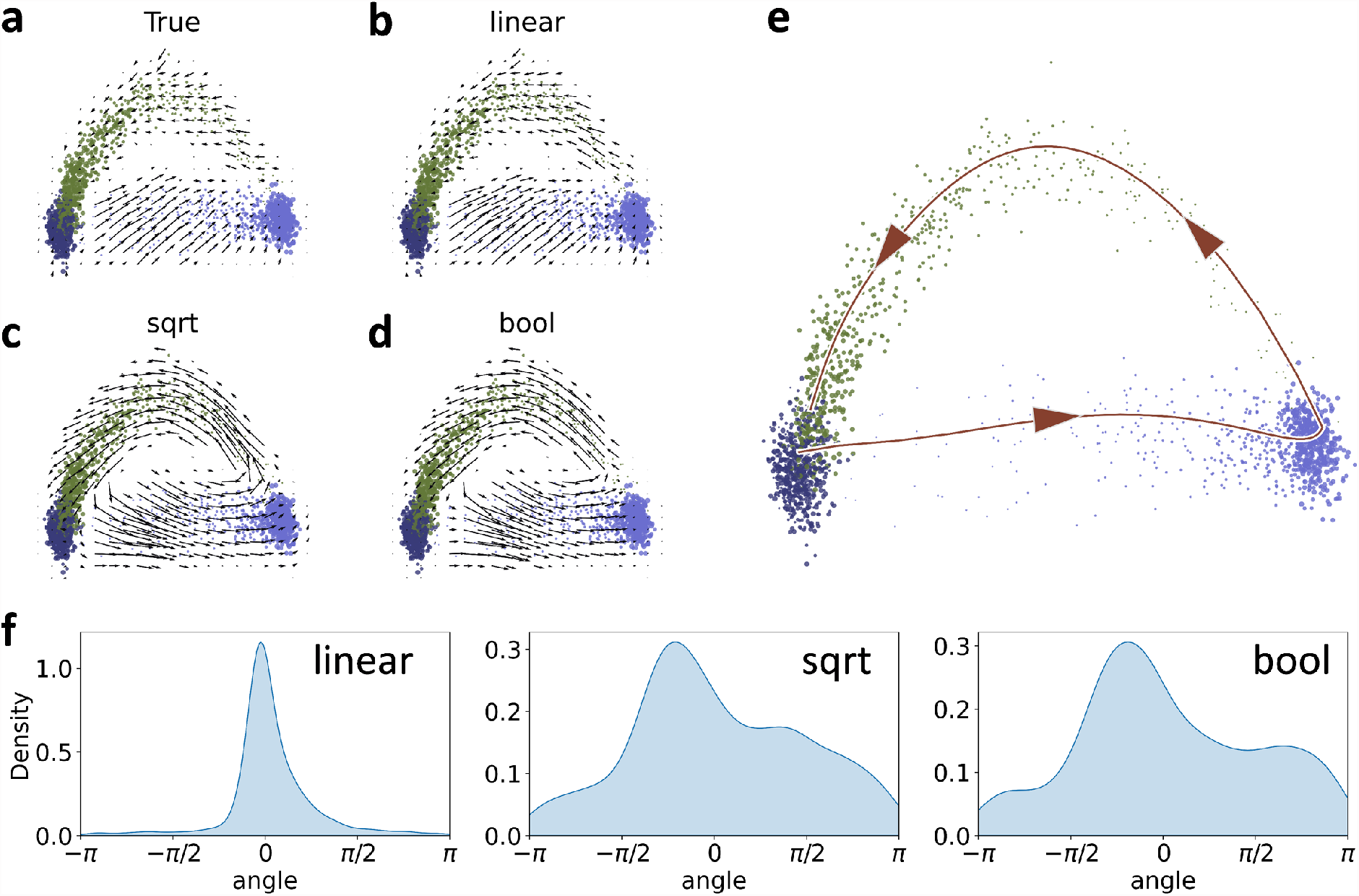
Performance of cell and velocity embeddings on simulated data, compared to ground truth velocity directions. **a**. Linear PCA embedding of ground truth velocities. **b**. Linear PCA embedding of inferred velocities. **c**. Nonlinear PCA embedding of inferred velocities. **d**. Nonlinear, Boolean PCA embedding of inferred velocities. **e**. Embedding of ground truth principal curve; trajectory directions displayed to guide the eye. **f**. Distribution of cell-specific angle deviations relative to ground truth velocity directions.

## 3 Logic and methodology

To understand the implications of the choices implemented in various RNA velocity workflows, we examined the procedures from a biophysics perspective, with a view towards understanding the mechanistic and statistical meaning of methods implemented. In this section, we broadly discuss potential challenges, problematic assumptions, and contradictory results. In the following section, we draw on lessons learned and propose a modeling approach of our own.

### 3.1 Pre-processing

As outlined in Section 2.1, several workflows are available for converting raw reads to molecule counts. These workflows largely follow the logic set out in the original implementation [1]; however, as pointed out by Soneson et al. [28], they produce different outputs from the same data. We reproduced their analysis on a broader selection of datasets (Section 6.9) in Figure S3, according to the procedure in Section 6.1. The performance was broadly consistent with the previous benchmarking and the description in Section 2.1: the methods agreed on the definition of “spliced” molecules, but different rules for the assignment of “unspliced” molecules led to discrepancies in counts. These discrepancies were particularly pronounced when comparing datasets one gene at a time, likely due to noise in tens of thousands of low-expressed genes (*ρ* by cell in Figure S3; cf. lower triangle of Fig. S10 in [28]).

However, a simple comparison between the software outputs obscures a far more fundamental challenge: the binary classification of transcripts as either spliced or unspliced is necessarily incomplete. The average human transcript has 4–7 introns [66], and a combinatorial number of potential transient and terminal isoforms. The vast majority of genes are alternatively spliced [67–69].

We can consider the hypothetical example of a nascent transcript with the structure *E*_1_*I*_1_*E*_2_*I*_2_*E*_3_, where *I*_*i*_ are introns and *E*_*i*_ are exons, as shown in Figure 4. If we place all UMIs with intronic reads into the “unspliced” category, we conflate the parent and intermediate transcripts. On the other hand, if we place all UMIs with splice junctions into the “spliced” category, we conflate the intermediate and terminal transcripts. Adding more complexity, some isoforms may retain introns through alternative splicing mechanisms; for example, the intermediate transcripts may be exported, translated, and degraded alongside the terminal.

The binary model is not large enough to include the diversity of possible splicing dynamics, but approximately holds under fairly restrictive conditions: the predominance of a single terminal isoform, as well as the existence of a single rate-limiting step in the splicing process. Previous work reports that minor isoforms are non-negligible [67, 69], differential isoform expression is physiologically significant [69–71], and intron retention in particular is implicated in regulation and pathology [72–75].

Splicing rate data are more challenging to obtain, but targeted experiments [76], genome-wide imaging [77], and our preliminary mechanistic investigations [78] suggest that selection and removal of individual introns is stochastic, but the overall splicing process has rather complex kinetics, not reducible to a single step.

### 3.2 RNA velocity biophysics

We will first inspect the complexity obscured by the simple schema given in Figure 3a. The velocity manuscripts use several distinct models for the transcription rate *α*(*t*). Furthermore, the amounts of molecular species U and S (previously denoted informally by *u* and *s*) have incompatible interpretations. The following models make fundamentally different claims about the data-generating process and imply fundamentally different inference procedures.

1. *α*(*t*) is piecewise constant over a finite time horizon; *u* and *s* are discrete (SN2 pp. 2-3, Fig. 1a-b of [1]).
2. *α*(*t*) is continuous and periodic; *u* and *s* are discrete (SN2 pp. 2-3, Fig. 1e of [1]).
3. *α, β*, and *γ* all smoothly vary over a finite time horizon according to an undisclosed function, with *α* exhibiting pulse behavior; *u* and *s* are discrete (SN2 Fig. 5 of [1]).
4. *α, β*, and *γ* all smoothly vary over a finite time horizon according to an arbitrary function; *u* and *s* are continuous (Fig. 3 of [22]).
5. *α*(*t*) is piecewise constant over a finite time horizon; *u* and *s* are continuous (Fig. 1 of [1], Methods of [3]).
6. *α*(*t*) is piecewise constant over a finite time horizon; *u* and *s* are continuous-valued but may contain discontinuities (Methods of [3]).
7. *α* is constant; *u* and *s* are continuous (Fig. 1b and SN1 pp. 1-2 of [1]). This formulation yields the reaction rate equation, and cannot produce the bimodal phase plots of interest.
8. *α* is constant; *u* and *s* are discrete (SN1 pp. 2-3 of [1]). This is the stochastic extension [79] of the previous model, and cannot produce the bimodal phase plots of interest, as explicitly shown on page 3 of SN1 in [1].

These discrepancies make a comprehensive analysis challenging. Models 7-8 do not contain differentiation dynamics. Certain models are contrived; models 3-4 propose transcription rate variation without motivating the specific form, and model 6 introduces nonphysical discontinuities. Model 2 alludes to limit cycles in stochastic systems under periodic driving, an intriguing phenomenon in its own right [80, 81], but not otherwise explored in the *scVelo* and *velocyto* publications. For the rest of this report, we focus on the discrete formulation (model 1) and its continuous analog (model 5).

For the discrete formulation, the RNA velocity *v* should be interpreted as the time derivative of the expectation of a random variable *S*_*t*_ that tracks the number of spliced RNA, conditional on the current state:

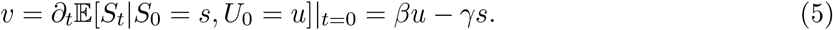

For the continuous formulation, it should be interpreted as the time derivative of the deterministic variable *s*_*t*_ that tracks the amount of spliced RNA, initialized at the current state:

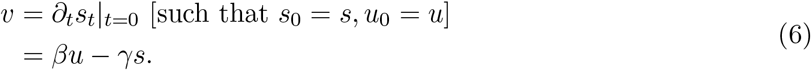

These formulations happen to be mathematically identical, which creates ambiguity. Nevertheless, both are legitimate, if narrow, statements about the near future of a process initialized at a state with *u* unspliced and *s* spliced molecules. The questions that arise immediately before, and immediately after, the velocity computation procedure, are (1) what generative model should be fit to obtain *β* and *γ* and (2) even with a *v*, how much use can one make of it?

### 3.3 Model definition

Continuous, deterministic models are fundamentally inappropriate in the low-copy number regime, which is predominant across the transcriptome [82–84]. Although continuous equations such as Equation 1 can represent the evolution of moments, they are insufficient for inference, as fitting average mRNA abundance amounts to invoking the central limit theorem for a very small, strictly positive quantity [85–89]. A comprehensive understanding of the stochastic noise model is necessary prior to making such approximations. Therefore, simulation methods that use a continuous model are immediately suspect [16]. We describe implementation-specific concerns in Sections 3.4 and 3.5.

The motivation behind a pulse model of transcriptional regulation is obscure. Although dynamic processes certainly have transiently expressed genes [90–92, 92, 93], it is far from clear that this model applies across the transcriptome, to thousands of potentially irrelevant genes. Indeed, it is not even coherent with genes showcased in the original report (Fig. 4d and Extended Data Fig. 8b of [1]): only ELAVL4 appears to show a symmetric pulse of expression. Finally, even when this model does apply, the assumption of constant splicing and degradation rates across the entire lineage is a potentially severe source of error, with no simple way to diagnose it [22].

Most problematic is that even the discrete model is incoherent with known mammalian transcriptional dynamics. If we suppose induction and repression periods are relatively long, as for a stationary, terminal, or unregulated cell population, we arrive at genome-wide constitutive transcription, in which the rate of RNA production is constant. This contradicts numerous sources that suggest transcriptional activity varies with time even in stationary cells [87, 94–101], and is effectively described by a telegraph model that stochastically switches between active and inactive states [102, 103].

Thus, we must impose basic consistency criteria. Using the models in Section 3.3 requires the assumption that stationary, homogeneous cell populations are Poisson-distributed. This assumption contradicts at least thirty years of evidence for widespread bursty transcription [95, 103]. We have obtained the answer to question (1) in Section 3.2: the model must be coherent with known biophysics, and provide a robust way to identify cases when its assumptions fail.

### 3.4 Count processing

Some standard properties of constitutive systems appear to at least qualitatively motivate gene filtering. Only genes with spliced–unspliced Pearson correlation above 0.05 are used for fitting parameters (as on p. 4 of SN2 in [1]); if the correlation is below this threshold, the gene is removed from procedure and presumed stationary. This is valid for the constitutive model, but inappropriate for broader model classes: for example, bursty transcription yields strictly positive correlations, making this statistic ineffective for identifying dynamics [78, 104].

Normalization relative to the cell’s molecular count is a standard feature of sequencing workflows [36, 105, 106], but reduces interpretability. Normalization converts absolute discrete count data to a proportion of the total cellular counts, ostensibly to account for the compositional nature of read-data data [107]. Several recent studies strongly discourage normalization of UMI-based counts [108, 109], although this perspective is not universal [110, 111]. It is clear that continuous-valued normalized data are incompatible with discrete mechanistic models. Moreover, the suitability of continuous models (such as Equation 1) is never explicitly justified, but merely assumed. Since normalization nonlinearly transforms the molecule distributions [65, 109] and introduces a coupling even between independent genes, the precise interpretation of single-gene ODE models is unclear.

Nearest-neighbor averaging is used to smooth the data after normalization. Though it effaces much of the stochastic noise to give an “averaged” trajectory, it introduces distortions of unknown magnitude. As discussed in Section 2.3, the imputation step does not have a consistent interpretation. The original report [1] defines it as “*k*NN pooling” in the manuscript and “imputation” in the documentation (and figure 17 of SN2), placing the emphasis on denoising. On the other hand, *scVelo* interprets the local average as an estimate of the expectations *μ*_*u*_(*t*), *μ*_*s*_(*t*). Neither approach appears to be justified by previous studies or benchmarking on ground truth, and both are circular as the neighborhood is computed based on the observed counts. A probabilistic analysis in Section S1.1 formalizes more deep-seated issues with using model-agnostic point estimates to “correct” data. Although these claims may hold and simply require more theoretical work to prove, our simulations in Section 4.3 strongly suggest they are invalid even in the best-case scenario: the phase portraits are smoothed out, but fail to capture the underlying dynamics in a way coherent with those claims.

To illustrate these problems, we performed a simple test of self-consistency, illustrated in Figure 5. We reprocessed the forebrain dataset (Fig. 4 of [1]) using the *velocyto* workflow, varying *k*, and investigated its effect on the appearance of the phase plots and the inferred parameters. As the neighborhood size was increased, the phase plot was distorted, with no apparent “optimal” choice of *k*.

### 3.5 Inference

Broadly speaking, *velocyto*-like moment estimates for *γ/β* are legitimate if the system has time to equilibrate (Section 4.2). However, moment-based estimation underperforms maximum likelihood estimation in general. The two approaches are in concordance under the highly restrictive assumptions of error normality and homoscedasticity. These assumptions are routinely violated in the low-copy number regime [85].

Regression on top and bottom quantiles inherits all of the issues of regression on the entire dataset, but compounds them by discarding a large fraction of data. Extremal quantile regression is otherwise a well-developed method [112–115], but it is generally applied to processes with nontrivial tail effects. For the quantile computation the filtering criterion is *ad hoc*, and not amenable to theoretical investigation. The order statistics of discrete distributions are notoriously challenging to compute [116–118], and even the simplest Poisson case exhibits complex trends [119]. In other words, the extrema themselves may be affected by noise, introducing more uncertainty into inference. Although the original article does perform some validation (SN2, Sec. 3 of [1]), it focuses on cell-specific velocities rather than parameter values, and only provides relative performance metrics rather than actual comparisons to simulated ground truth.

Even without testing the inference procedures against simulations, we can characterize their performance in terms of internal controls. As we demonstrate in Figure 5, the inferred *γ/β* values were unstable under varying *k*: the *velocyto* parameter inference procedure was highly sensitive to a user-defined neighborhood hyperparameter. On the other hand, using a simple ratio of the means (as in the first column and the *k* = 0 case in the fifth column of Figure 5) produced biases [1].

Interestingly, the fraction of cells predicted to be upregulated is qualitatively more stable, suggesting that the inference step is best understood as an *ad hoc* binary classifier, rather than a quantitative descriptor of system state. Given the stability of this classifier, as well as our preliminary discussion of similar results in the context of validating *protaccel* [4], we used this binary classifier as a benchmark in Sections 3.6 and 4.10.

Regression of the piecewise deterministic “dynamical” model in *scVelo* asserts the imputed counts have normal noise with equal residuals for spliced and unspliced species, once again implausible in the low-copy number regime. More fundamentally, it fails to preserve gene-gene coherence. If a cell is predicted to lie at the beginning of a trajectory for one gene, this estimate does not inform fitting for any other gene. The “dynamical” model appears to address this discrepancy in a *post hoc* fashion. First, the algorithm identifies putative “root cells,” which are themselves computed from the velocity embedding. Then, the disparate gene-specific times are aggregated into one by computing a quantile near the median. This procedure presupposes that the velocity graph is self-consistent and physically meaningful, and that the point estimate of pseudotime is sufficient, but does not mathematically prove these points or test them by simulation.

### 3.6 Embedding

After inference and evaluation of Δ*s* for every cell and gene, the array is converted to an embedding-specific representation. In the single-cell sequencing field, low-dimensional projections are more than a visualization method: they are ubiquitous tools for inference and discovery. Transcriptomics workflows convert large data arrays to human-parsable visuals; these visuals are then used to explore gene expression and validate cell type relationships, under the assumption that they represent the underlying data well enough to draw conclusions. However, the embedding procedures involve several distortive steps, which should be recognized and questioned.

For such visuals, the goal is to recapitulate local and global cell-cell relationships. However, accurately representing desired properties such as pairwise relationships between many points is inherently difficult, requiring dimensions several orders of magnitudes larger than two to faithfully represent the data [120]. Thus distortion of cell-cell relationships is naturally induced in two-dimensional embeddings, and grows as 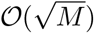 for *M* cells [120, 121]. Both linear (PCA) and nonlinear (*t*-SNE/UMAP) methods exhibit these distortions, and warp existing cell-cell relationships or suggest new ones not present in the underlying data [120, 122]. Tuning algorithm parameters can slightly improve some distortion metrics, though often at the expense of others [123]. Essentially, nonlinear embeddings utilize sensitive hyperparameters that can be tuned, but do not provide well-defined criteria for an “optimal” choice [122, 123]. Using visualizations for discovery thus risks confirmation bias.

The velocity algorithms present a particularly natural criterion for quantifying the embedding distortion. The nonlinear embedding procedure generates weights for vectors defined with reference to embedding neighbors. Therefore, we can reasonably investigate the effect of the embedding on the neighborhood definitions. In other words, if the velocity arrows quantify the probability of transitioning to a set of cells, what relationship does this set have to the set of neighbors in the pre-embedded data?

This relationship is conventionally [122] quantified by the Jaccard distance, defined by the following formula:

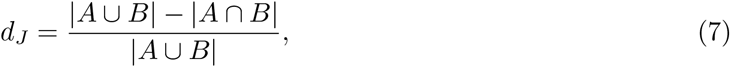

which reports the normalized overlap between the original sets of cell neighbors (A) and embedded cell sets (B). This dissimilarity metric ranges from 0 to 1, where 1 (or 100%) denotes completely non-overlapping sets. We applied standard steps of dimensionality reduction and smoothing to the forebrain dataset (Fig. 4 of [1]) and computed their effect on the neighborhoods (taken to be *k* = 150 for consistency with the velocity embedding process). We report the Jaccard distance distributions in Figure 6, and observe the gradual degradation of neighborhoods. On average, moving from the ambient high-dimensional space to a two-dimensional representation induced a *d*_*J*_ of 70-75%. Therefore, cell embedding substantially distorts precisely the local structure relevant to velocity embedding.

The two-dimensional arrows in Figure 1 combine three sources of error: the intrinsic information loss of low-dimensional projections, the instabilities in upstream processing and inference, and any additional error incurred by the nonlinear procedure outlined in Section 2.4. The softmax kernel-based procedure exhibits an inherent tension that merits closer inspection. On one hand, it is explicitly designed [1] to mitigate error incurred by cell- and gene-specific noise by performing several steps of pooling and smoothing. On the other hand, it is technically questionable: if we assume differentiation processes are largely governed by a small set of “marker genes,” pooling them with thousands of non-marker genes amounts to hoping that variation in an orthogonal data-generating process cancels out well enough to recapture latent dynamics. Certain processes may involve the modulation of large sets of genes, e.g., if expression overall increases over the transcriptome. However, the velocity workflows are intrinsically unable to identify such trends, as they use normalized data. As we demonstrate later, a model with no latent dynamics at all (Figure S6) can generate apparent signal in the embedded space, illustrating the dangers of relying on error cancellation. When multiple data-generating processes are present, naïve aggregation risks obscuring rather than revealing signal.

Aside from this high-level inconsistency, other problems emerge upon investigating the embedding procedure closer, even prior to performing any numerical controls. The nonlinear embedding approach introduced by La Manno et al. (Equation 4) is highly hyperparametrized, not motivated by any previous theory, has no physical interpretation, and does not appear to have been formally validated against linear or simulated ground truth. Just as with the cell embedding, the procedure is dependent on an arbitrary number of nearest neighbors and velocity transformation functions, with no clearly optimal choices. These hyperparameters can be tuned to correct for such instabilities, potentially resulting in overfitting to a pre-determined hypothesis. Since the procedure has no physical basis, potential false discoveries are challenging to diagnose. Furthermore, it reduces the limited biophysical interpretability of the result. The velocity derivation is model-informed and, as discussed in Sections 4.2 and 4.4, can be informally viewed as an approximation under several strong assumptions about the process biophysics. The embedding, on the other hand, is *ad hoc* and can only degrade the information content.

A final theoretical point remains before we can begin quantitatively validating the embeddings: as suggested by Equation 2, and discussed in Section 2.3, the *velocyto* gene-specific *v*_*j*_ have different units. Therefore, the aggregation in Equation 4 is questionable. The standard *velocyto* workflow assumes that the splicing rates are close enough to neglect differences, which appears to contradict other results reported in the same paper (Extended Data Fig. 2f of [1]).

To bypass this limitation in a self-consistent way, we implemented a “Boolean” or binary measure of velocity, as motivated by validation in the original manuscript (Sec. 3 in SN2 of [1]), introduced in the context of validating *protaccel* [4], and implied by resampling *β* values from a uniform distribution in an investigation of latent landscapes (p. 3 in Supplementary Methods of [13]). Essentially, instead of computing transition probabilities based on the velocity values, we computed them based on signs, bypassing the unit inconsistency. The algorithm used to produce this embedding is described in Section 6.8.

The Boolean procedure offers a natural internal benchmark. If this approach largely recapitulates findings from the standard methods, the embedding process serves as an information bottleneck: the inference procedure performs as well as a binary classifier, and the complexities of the dynamics are effaced by embedding. We used this approach as a trivial baseline and compared it to the standard suite of variance-stabilizing transformations implemented in *velocyto*. In addition, we tested the effects of neighborhood sizes, in the vein of the stability analysis performed in the original manuscript (Sec. 11 in SN2 of [1]). In Figure S4, we plot the distributions of angle deviations between a linear baseline, obtained by projecting the extrapolated cell state and computing *E*(*s*_*i*_ +Δ*s*_*i*_) − *E*(*s*_*i*_) using PCA, and the nonlinear velocity embedding. This control has not previously been investigated in any detail, but seems key to the claim that the nonlinear velocity embedding is meaningful: intuitively, we expect it to recapitulate the simplest baseline, at least on average. To avoid any confusion, we reiterate that the linear embedding is given by Equation 3, and not the identity nonlinear embedding implemented in *velocyto* (i.e., ϱ(*x*) = *x* in SN1, pp. 9-10 of [1]).

The angle deviations in arrow directions were all severely biased relative to the linear baseline. The different normalization methods were distortive to approximately the same degree. The performance of the Boolean embedding, which discards nearly all of the quantitative information, was nearly identical to the built-in methods, which suggests that the choice of normalization methods is a red herring: quantitative velocity magnitudes have little effect on the embedding quality. This is consistent with previous investigations (cf. Fig. S52 in [4]). On the other hand, the neighborhood sizes did not appear to matter much, at least over the modest range explored here (in contrast to Sec. 11 in SN2 of [1]). Therefore, the directions reported in embeddings were unrepresentative of the actual velocity magnitudes in high-dimensional space, as well as severely distorted relative to the linear projection. These discrepancies are a potential cause for concern. Observing the qualitative similarity of Figs. 2d and 2h in the original report [1], the reasonable performance of the linear extrapolation in *t*-SNE in its supplement (SN2 Fig. 9a of [1]), as well as the cell cycle dynamics explored with the linear embedding in *Revelio* [11], a casual reading of the RNA velocity literature would suppose these embeddings to be largely interchangeable.

Finally, we visualized the aggregated velocity vectors in Figure 7 to assess the local and global structures. This visualization served as both an internal and an external control. The internal control demonstrated the local structure and the stability of the velocity-specific methods, i.e., the actual directions of the arrows on the grid. We compared the conventional nonlinear projection to the Boolean method, as well as the linear embedding. The external control concerned the global structure, which can be analyzed in light of known physiological relationships: radial glia differentiate through neuroblasts into neurons [1]. If this global relationship is not captured by the embedding, the inferred trajectories are *a priori* physically uninterpretable, in a way that is particularly challenging to diagnose.

In the PCA embedding, the global structure was retained and the arrows were fairly robust, even when the non-quantitative Boolean method was used. However, the various projection options suggested drastically different relationships between the cell types, with PCA presenting more continuous representations of cell relationships faithful to ground truth, and UMAP and *t*-SNE presenting more local images, with distinct and discrete clusters of cells. Clearly, if the relationship between progenitor and descendant is lost, the velocity workflow cannot infer it. The *t*-SNE and UMAP parameters can be adjusted by the user; however, adding a new set of tuning steps and optimizations provides an opportunity for confirmation bias to overrule the data.

### 3.7 Summary

The standard RNA velocity framework presupposes that the evolution of every gene’s transcriptional activity throughout a differentiation or cycling process can be described by a continuous model with a single upregulation event and a single downregulation event. It proceeds to normalize and smooth the data until the rough edges of single-molecule noise are filed off, fitting the continuous model assuming Gaussian residuals.

In the process, the stochastic dynamics that predominate in the low-copy number regime, and characterize nearly all of mammalian transcription, are lost and cannot be recovered. Although parameters can be fit, they are distorted to an unknown extent, due to a combination of data transformation, suboptimal inference, and unit incompatibilities. The gene-specific components of velocity are underspecified due to their direct dependence on the imputation neighborhood and splicing timescale. In *scVelo*, parameters are estimated under a highly restrictive model, yet applied to make broad claims about complex topologies. In *velocyto*, only the sign of the velocity is physically interpretable. It can still be used to calculate low-dimensional directions, and this binary velocity embedding is seemingly as good as any other, suggesting the other methods lose information. However, the embedding process itself is not based on biophysics, and is not guaranteed to be stable or robust. Fortunately, the natural match between stochastic models and UMI-aided molecule counting offers the hope for quantitative and interpretable RNA velocity.

## 4 Prospects and solutions

*Is there no balm in Gilead?* Given the foundational issues we have raised, how can the RNA velocity framework be reformulated to provide meaningful, biophysically interpretable insights? We propose that discrete Markov modeling can directly and naturally address the fundamental issues. In particular, transient and stationary physiological models can be defined and solved via the chemical master equation (CME), which describes the time evolution of a discrete stochastic process. Since the “noise” is the data of interest, in such an approach smoothing is not required. Rather, technical and extrinsic noise sources can be treated as stochastic processes in their own right, and explicit modeling of them can improve the understanding of batch and heterogeneity effects. Finally, within this framework, parameters can be inferred using standard and well-developed statistical machinery.

### 4.1 Pre-processing

The diversity of potential intermediate and terminal transcripts suggests that simplistic splicing models are inadequate for physiologically faithful descriptions of transcription dynamics. What is needed is a treatment of the types of transcripts listed in Section 3.1 as distinct species. This approach immediately leads to several significant challenges, relating to quantification, biophysics, and identifiability.

Transient, low-abundance intermediate transcripts are substantially less characterized than coding isoforms. Some data are available from fluorescence transcriptomics with intron-targeted probes [46], but such imaging is impractical on a genome-wide scale. Unfortunately, the references and computational infrastructure necessary to identify intermediate transcripts do not yet exist.

Even if intermediate isoforms could be perfectly quantified, single-cell RNA-seq data do not generally contain enough information to identify the order of intron splicing. The problem of splicing network inference has been examined; however, experimental approaches [124, 125] are challenging to scale, whereas computational approaches [126] do not generally have enough information to resolve ambiguities.

Furthermore, even with complete annotations and a well-characterized splicing graph at hand, large-scale short-read sequencing cannot fully resolve transcripts. This limitation gives rise to a challenging inference problem. For example, if transcripts *A* := *E*_1_*I*_1_*E*_2_*I*_2_*E*_3_ and *B* := *E*_1_*E*_2_*I*_2_*E*_3_ are indistinguishable whenever only the 3’ end of each molecule is sequenced, it is necessary to fit parameters through the random variable *X*_*A*_ + *X*_*B*_, i.e., from aggregated data. The functional form of this random variable is not analytically tractable in general.

We have described a preliminary method that can partially bypass these problems [78]. Sequencing “long” reads, which at this time is possible with technologies such as Oxford Nanopore [69], or sequencing of “full-length” libraries produced with methods such as Smart-seq3 [127], enhances identifiability and facilitates the construction of new annotations based on presence or absence of intron sequences. Finally, even though additional data are required to specify entire splicing networks, sequencing data are sufficient to constrain parts of these networks; for example, if two transcripts differ by one intron, the longer one cannot possibly be generated from the shorter.

Defining more species leads to inferential challenges in downstream analysis. Even if sequencing data are available, their relationship to the biological counts is nontrivial: some intermediate transcripts may not be observable using certain technologies because they do not contain sequences necessary to initiate reverse priming, whereas others may be over-represented in the data because they contain many. In a preliminary investigation [128], which adopts the binary categories of “spliced” and “unspliced” defined in the original RNA velocity publication, we found that unspliced molecules originating from long genes are overrepresented in short-read sequencing datasets. This suggests that multiple priming occurs at intronic poly(A) sequences. To “regress out” this effect, a simple length-based proxy for the number of poly(A) stretches can be used, but a more granular description would require a sequence-based kinetic model for each intermediate transcript’s capture rate.

### 4.2 Occupation measures provide a theoretical framework for scRNA-seq

The data simulated in the exposition of RNA velocity [1] comes from a particular set of what are called Markov chain occupation measures. As an illustration of what this means, we consider the simplest, univariate model of transcription, a classical birth-death process:

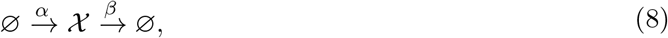

where *α* is a constant transcription rate and *β* is a constant efflux rate. Depending on the system, *β* may have various biophysical interpretations, such as splicing, degradation, or export from the nucleus [78, 104].

Formally, the exact solution to this system is given by the chemical master equation (CME), an infinite series of coupled ordinary differential equations that describe the flux of probability between microstates *x*, which specify the integer abundance of 𝒳, defined on ℕ_0_:

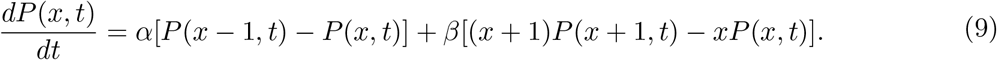

This equation encodes a full characterization of the system: transcription is zeroth-order, efflux is first-order, and the dynamics are memoryless, i.e., depend only on the state at *t*. We define the quantity *y*, namely the solution to the underlying reaction rate equation that governs the average of the 𝒳 copy number distribution:

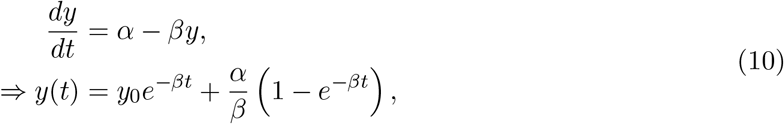

where *y*_0_ is the average at time *t* = 0. To simplify the analysis, we assume that the initial condition is Poisson-distributed. Per classical results [79], the distribution of counts *P* (*x, t*) is described by a Poisson law for all *t*, and converges to *Poisson*(*α/β*) as *t* → ∞. Quantitatively, the time-dependent distribution is given by *P* (*x, t*) ∼ *Poisson*(*y*(*t*)):

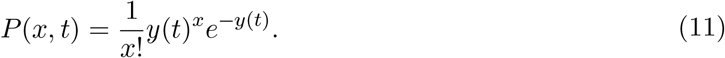

However, this is not the correct model class for distributions observed in scRNA-seq datasets. To appreciate these subtleties, we delve into and interrogate assumptions that underpin the use of such distributions.

The sequencing process does not “know” anything about the transcriptional dynamics and their time *t*. This stands in contrast to transcriptomics performed *in vitro*, with a physically meaningful experiment start time. For example, in many standard protocols, a stimulus is applied to the cells at time *t* = 0, and populations of cells are chemically fixed and profiled at subsequent time points, potentially up to a nominal equilibrium state [2, 33, 85, 129]. However, if there is no experimentally imposed timescale, and we adopt the standard assumption that cell dynamics are mutually independent, the process time decouples from experiment time. Although cells are sampled simultaneously, their process times *t* are draws from a random variable that must be defined.

Formalizing this framework requires introducing the notion of occupation measures. Considering a single cell, we designate its process time *t* as a latent pseudotime. Our definition of this term is at odds with standard use. In brief, “pseudotime” conventionally denotes a one-dimensional coordinate along a putative cell trajectory, which parameterizes a principal curve in a space based on RNA counts [36]. In our case, we use this term to denote a real time coordinate, which governs the process “clock.” This difference is fundamental. The Markov chain pseudotime is physically interpretable as the progress of a process that induces the observations in expression space. Conversely, the expression pseudotime is purely phenomenological, and we are unaware of any trajectory inference methods that explicitly parameterize the underlying stochastic model using the CME; instead, all available implementations appear to use isotropic or continuous noise models [36, 106, 130–138]. As we emphasize in Section 3.3, these models are inappropriate for low-abundance molecular species.

By construction, cell trajectories are observed at times *t* ∈ ℝ. This requires introducing a sampling distribution *f* (*t*), which describes the probability of observing a cell at a particular underlying pseudotime. Therefore, the probability of observing *x* molecules of X in the constitutive case takes the following form:

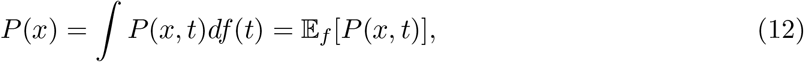

i.e., the expectation of *P* (*x, t*) under the sampling law. *P* (*x*) is called the occupation measure of the process, and reports the probability that a trajectory is observed to be in state *x*, a slight generalization of the usual definition [139–141].

Next, we must encode the assumption that cell observations are desynchronized from the sequencing process and each other. This assumption leads us to a choice consistent with the previous reports [1, 22], namely *df* = *T* ^−1^*dt*, where [0, *T*] is the pseudotime interval observable by the sequencing process. This constrains the probability of observing state *x* to be the actual fraction of time the system spends in that state. Then, we take *T* → ∞, yielding

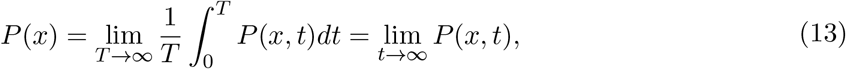

which is a statement of the ergodic theorem [142]. Under mild conditions, this theorem guarantees that samples from unsynchronized trajectories converge to the same distribution as the far more tractable ensembles of synchronized trajectories.

With this discussion, we have clarified that the application of stationary distributions lim_*t*→∞_ *P* (*x, t*) to describe ostensibly unsynchronized cells naturally emerges from assumptions about biophysics and the nature of the sampling process. However, these assumptions may be violated; for example, RNA velocity describes molecules sampled from a transient process. This distinction is key: limits such as lim_*t*→∞_ *P* (*x, t*) may not even exist, and we expect to capture only a portion of the trajectory. A rigorous probabilistic model must treat the occupation measure directly, which remains valid without those assumptions. Formally, this amounts to relaxing the assumption of desynchronization: the sequencing process is time-localized to a particular interval of the underlying biological process.

To stay consistent, we continue using the sampling law *df* = *T* ^−1^*dt* on [0, *T*], but infinite supports are valid so long as they decay rapidly enough to be integrable. As scRNA-seq data are atemporal, this time coordinate is unitless and cannot be assigned a scale without prior information, so *T* can be defined arbitrarily without loss of generality.

The occupation measure of the birth-death process takes the following form:

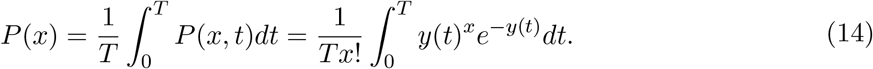

This integral can be solved exactly; however, this solution does not easily generalize to more complex systems. Instead, we can consider the probability-generating function (PGF), which also takes a remarkably simple form:

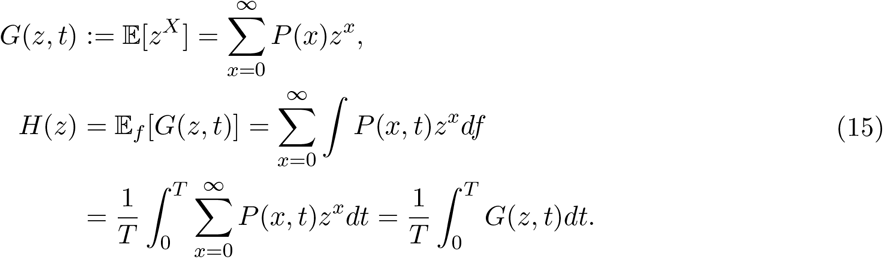

By linearity, the generating function *H*(*z*) of the occupation measure is simply the expectation of the generating function *G*(*z, t*) of the original process with respect to the sampling measure *f*. From standard properties of the birth-death process, this yields:

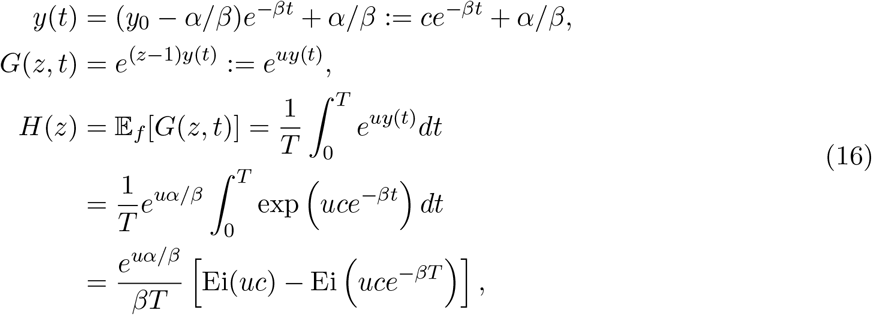

where *Ei* is the exponential integral [143]. This is the solution to the system of interest. Although straightforward to evaluate, it does not appear to belong to any well-known parametric family. At this point, several properties relevant to the standard definition of velocity stand out:

Extending the solution to multiple species merely requires defining the correct multivariate *G*(*z, t*). This is tractable for all directed acyclic graphs of splicing [78]. However, the broader set of models does not have an analytical solution, even in terms of the exponential integral, because it requires integrals of the following intractable form:

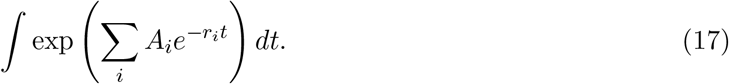

Extending the solution to piecewise constant transcription rates, as in Figure 3, or to any piecewise constant parameter values, only requires computing the appropriate *y*(*t*). This is straightforward to do by using the initial conditions on disjoint intervals, and generalizes to multi-species reaction systems.

The model is broad enough to describe cell type distinctions and cell fate stochasticity simply by defining a discrete mixture model over transcription rate trajectories. For example, if a cell can choose to enter cell fate *A* with probability *w*_*A*_ or cell fate *B* with probability 1 − *w*_*A*_, the overall generating function takes the following form:

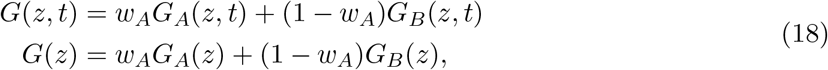

where each branch’s generating function is induced by distinct driving functions, *α*_*A*_(*t*) ≠ *α*_*B*_(*t*).

Finally, as discussed in Section 3.3, the constitutive model is problematic. Fortunately, the occupation measure formulation is robust enough to deal with more general systems. For example, we can consider the classical example of a bursty system, extended to include deterministic transcriptional modulation. The following chemical schema represents the system dynamics:

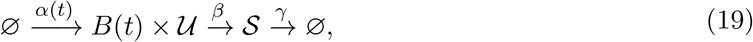

where *α*(*t*) is the burst frequency and *B* is the burst size, a random variable drawn from a geometric distribution on ℕ_0_ with expectation *b*(*t*). The log PGF of the system takes the following form:

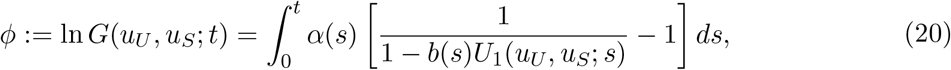

where *u*_*U*_ and *u*_*S*_ are generating function arguments and *U*_1_ is a tractable exponential sum [104]. By linearity, the generating function of the occupation measure takes the simple form *G*(*u*_*U*_, *u*_*S*_) = 𝔼_*f*_ [*G*(*u*_*U*_, *u*_*S*_; *t*)]. Just like the underlying generating function, this expression cannot be solved analytically and requires quadrature. For example, for *f* defined over a finite time horizon, it is straightforward to construct a discrete grid of *t*_*i*_ ∈ [0, *T*], evaluate the integrand at these times, and apply the trapezoidal rule to estimate the iterated integral through matrix-vector operations. With this mathematical formulation, we can elaborate on the answer to the first question in Section 3.2: the class of models with bursty transcription is consistent with known biophysics.

### 4.3 Count processing

Selecting and solving a model allows for a quantitative comparison of the predictions of the inference process to a meaningful baseline. Consider the simplest analytically tractable model, where *α*(*t*) is piecewise constant over a finite time horizon, with its dynamics given by a single positive or negative pulse:

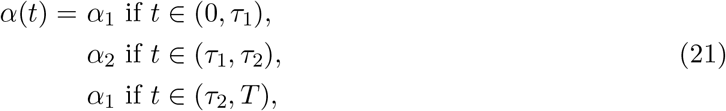

and *α*_1_ ≠ *α*_2_. Different genes have different *α*, but synchronization is enforced enforced: all genes switch at identical times *τ*_1_, *τ*_2_. Physically, this description corresponds to a perturbation being applied to, then removed from the cells, resulting in the modulation shown in Figure 8a. We simulated this model using the procedure outlined in Section 6.2.

We investigated this class of phenomenological models for transcription variation as opposed to fully mechanistic descriptions (e.g., *dyngen* [15]) for three reasons. First, they provide the best-case baseline scenario for the validation of RNA velocity, as the inference procedure is predicated on this model. For example, Equation 21 is the pulse stimulus model proposed by La Manno et al. [1]. Second, they offer the analytical solutions discussed in Section 4.2, whereas more complicated schema do not. Finally, mechanistic models of regulation are underdetermined relative to scRNA-seq data, as they rely on signal transfer through proteins. We anticipate that comprehensive future models will introduce further mechanistic details, as in the causal schema proposed in Figure 4 of [22]. However, here we abstract away the specific details of how this perturbation is effected, and focus on its effects on the observable transcriptome.

Predictably, the marginal distributions of the simulated data were bimodal, and matched the analytical solutions for the occupation measure (Figure 8b). The count processing workflow distorted the data in complex, nonlinear ways. Compared to the raw data, imputation did lower dispersion (Figure 8c). However, we did not find support for treating the imputed data as merely *μ*_*u*_(*t*) and *μ*_*s*_(*t*) with a Gaussian perturbation. For any particular gene, the relationship between the two was nontrivial and exhibited biases. The sample gene demonstrated in the figure produced an approximately smooth trace that did not resemble the true process average. At best, the imputed estimate appeared to be unbiased over the entire dataset, and tended to fall within of factor of ten of the true mean (Figure 8d i-ii). Interpreting the local observed variance as the true process moment *σ*^2^ is even less appropriate: the relationship between local and true variance exhibited unintuitive, nonlinear effects (Figure 8d iii-iv).

Normalization for the total number of UMI counts per cell is standard, and La Manno et al. do use it in their validation of *velocyto* (p. 6 in SN2 of [1]). On the other hand, it could be argued that such scaling is an *ad hoc* procedure intended to eliminate underlying variation in UMI counts due to differences in which genes are expressed in cells, or technical noise. As such, it may be inappropriate to apply it to simulated datasets that do not have these phenomena. We repeated the analysis without scaling counts by total molecule counts per cell in Figure S5. This means that raw counts were pooled, using the *k* nearest neighbors obtained from PCA, which was itself computed from the log-transformed raw, spliced counts. Qualitatively, the gene-specific imputed trajectories did not exhibit the severe biases of Figure 8. However, they still produced errors in the transient regimes of interest, and scale- and species-dependent errors elsewhere, violating the assumptions of the *scVelo* “dynamical” inference procedure (as outlined in Section 3.5). Furthermore, the relationship between true moments and pooled moments was nearly identical in Figure 8d and Figure S5d.

We can conclude that even the best-case scenario, with matching model assumptions and no technical noise, does not justify using imputed data in place of the process average: the imputation procedure is circular, and can give rise to biases (as in Section S1.1). Although these biases may cancel on average, this cannot be relied on for any particular gene. Therefore, instead of smoothing, which is fundamentally unstable and challenging to validate, we recommend explicitly constructing and solving error models (as in [128]). Such an approach provides insights into the system biophysics and enables the quantification of uncertainty through standard statistical methods.

### 4.4 Inference from occupation measure data

With the groundwork outlined in Sections 4.2 and 4.3, one can begin to tackle the problem of inference from sequencing data. We start by carefully inspecting the models set up in the velocity publications, enumerating their assumptions about the transcriptional processes, and writing down their formal solutions without making any further approximations. In general, the resulting problems are intractable. However, it is instructive to write down the exact forms, as they clarify the origin of the complexity and can point to solution strategies.

First, we define the global structure of the transient system, which represents the parameters shared between different genes. This global structure is encoded in the vector *θ*_*G*_, which we assume to be finite-dimensional. For example, if the differentiation process is linear and deterministic, with no branching, with *K* distinct cell types, *θ*_*G*_ gives the times *τ*_*k*_ of the cell type transitions (where *τ*_0_ := 0 and *τ*_*K*_ := *T*, which we set to 1 with no loss of generality). On the other hand, if it is non-deterministic, with diverging cell fates, *θ*_*G*_ can encode topologies and fates’ probabilities (such as *w*_*A*_ from Equation 18). Conversely, a unipotent differentiation trajectory can be encoded by setting *w*_*A*_ = 1, i.e., simpler topologies are degenerate cases of more complex topologies. Finally, this vector may also encode non-biological phenomena, such as batch-specific technical noise parameters [128] and the specific form of the generalized occupation measure *f*.

In addition to *θ*_*G*_, the system also involves vectors *θ*_*j*_, *j* ∈ {1, …, *N*}, where *j* ranges over the genes. The *θ*_*j*_ are gene-specific parameter vectors that parameterize physiological processes, such as transcriptional bursting, splicing, and degradation, as well as initial conditions. For simplicity, we make two crucial assumptions, consistent with the previous velocity publications. First, we suppose the transcriptional parameters are piecewise constant throughout the process, and all other parameters are strictly constant. Second, we suppose that different genes’ transcription and processing reactions are statistically independent. We define the full parameter set as Θ := {*θ*_1_, …, *θ*_*N*_, *θ*_*G*_}. For completeness, we note that in previous publications the cell type transition times are gene-specific parameters, i.e., *τ*_*jk*_ ∈ *θ*_*j*_, but since we assume that all genes switch at identical times they are global parameters, i.e., *τ*_*k*_ ∈ *θ*_*G*_, in the model we propose here.

Finally, we formalize the data variables. The full dataset is given by the data matrix 𝒟, with cell-specific arrays 𝒟_*i*_ and gene-specific arrays 𝒟_*j*_. Thus, 𝒟_*ij*_ reports the spliced and unspliced counts for gene *j* in cell *i*. Under this model and the assumption of gene independence, the following equation gives the likelihood of a particular cell’s observation:

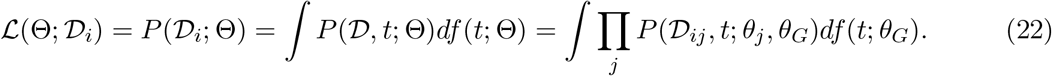

The assumption of cell independence suggest the following total likelihood:

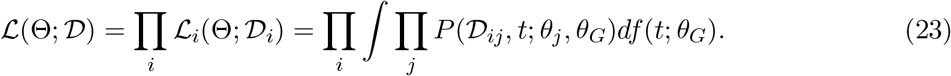

To fully characterize this system under the foregoing model assumptions, we need to optimize the likelihood with respect to the parameters. This is generally intractable; even evaluating Equation 23 can be non-trivial. However, we can make a series of simplifying approximations.

### 4.5 A combinatorial optimization approach to inference

Ultimately, we would like to understand the behavior of Equation 23 across the entire domain of parameters, compute the maximum likelihood estimate (MLE) of Θ, and characterize its stability by calculating its confidence region. However, this is not yet feasible. Therefore, we restrict the discussion to writing down a formula for the MLE that can be treated using standard algorithms.

Our strategy is to exploit the “latent” distribution of cell-specific process times, condition on this distribution, and find an approximate MLE. Assuming that cells are observed uniformly across pseudotime, i.e., *df* = *dt*, the latent time of each cell *i* is given by *t*_*i*_ ∈ (0, 1). These times are almost surely distinct, and induce a cell ranking in order of increasing pseudotime. This ranking is unknown and has to be inferred from the data.

Assume, for the moment, that the ranking is known and given by *σ*, a permutation of the *M* cell indices {1, 2, …, *M* − 1, *M*} corresponding to their pseudotime order statistics. Given a cell’s order statistic *σ*_*i*_, we can use *f* to estimate its latent time *t*_*i*_, which is distributed according to a rather complex multivariate Beta distribution [144]: even if each cell’s rank in the total order is known, we have to account for the uncertainty in pseudotime. However, we can exploit the fact that this uncertainty decreases as the number of cells grows. The marginal order statistics are distributed according to *Beta*(*σ*_*i*_, *M* + 1 − *σ*_*i*_), a random variable with law 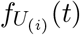 and the following variance:

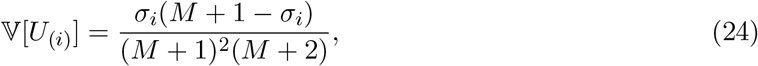

which tends to zero with the rate *M* ^−1^. Therefore, we find that 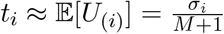.

Thus, if we have enough cells, and know their pseudotime ordering, we can exploit the fact that 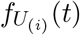 converges to a Dirac delta functional:

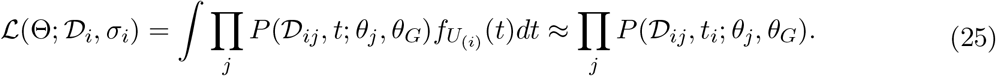

This amounts to using a plug-in point estimate to compute the likelihood of a cell’s data. The same approach can be applied to each cell in turn:

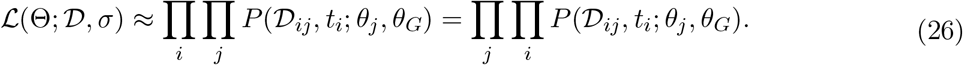

However, optimizing this quantity, even when conditioning on *σ*, is impractical because it requires simultaneously optimizing (potentially thousands of) gene-specific parameters along with the (relatively few) global parameters. The interchange of product operations is helpful because it allows us to write down a more tractable loss function for the MLE, which exploits the conditional separability of *θ*_*j*_:

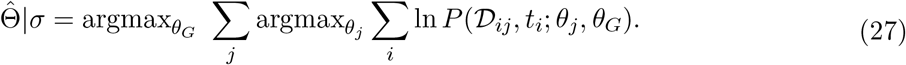

Optimizing this quantity over *σ* guarantees to return the global MLE:

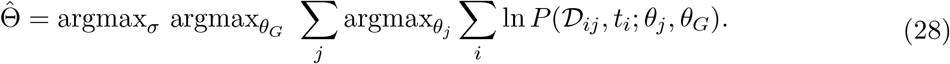

The parameters *θ*_1_, …, *θ*_*N*_, *θ*_*G*_ can be found by standard continuous optimization methods, but estimating *σ* requires a combinatorial optimization, namely finding an optimal traversal path between cells. In other words, even this approximate approach requires solving the problem of pseudo-time inference, which produces a one-dimensional ordering of cells [145]. However, unlike standard pseudotime inference, which describes a set of purely phenomenological relationships informed by proximity between cell expression states, the current theoretical framework endows the solution with a concrete biological interpretation, which is informed by a specific microscopic model of transcription.

The trajectory inference literature treats this class of problems by graph traversal algorithms, generally by constructing a minimum spanning tree or an optimal traversal on the clusters or individual cells [15, 137]. Under fairly severe modeling assumptions, which generally rely on error isotropy, the optimal traversal of cell states reduces to the traveling salesman problem (TSP) via the Hamiltonian path problem with some minor differences [130, 146, 147]. The current approach is considerably more complicated, because the weights between the “nodes”, i.e., the observed cells, cannot be generally written down in closed form, and require optimization for every *σ*.

Nevertheless, the specific form of the required combinatorial optimization has several useful implications for inference. It is possible to subsample or filter the data to obtain rough estimates of the parameters by sampling a subset of genes. This facilitates the estimation of *θ*_*G*_, which can be reused for an entire dataset. If only a fraction of genes are systematically modulated across a trajectory, technical noise parameters within *θ*_*G*_ can be estimated from the far more easily tractable fits to the stationary genes. Furthermore, sampling a subset of cells enables the construction of approximate *σ* and estimation of *θ*_*j*_. The validity of such approximations can be assessed with relatively simple controls. For example, if a best-fit “trajectory” over cells *σ* is as good as a random or inverted permutation, the transient model is likely overfit. Finally, existing trajectory inference methods can be exploited to obtain an ordering *σ* for the purpose of testing whether it can give results consistent with the stochastic model by calculating the optimal parameters in Equation 27, plugging them into the appropriate CME, and comparing the process occupation measure to the true molecule distributions.

It is plausible that many standard trajectory inference methods can be represented as approximations to the exact solution under particular assumptions about the form of biological model and priors imposed on the trajectory structure. However, a complete discussion of the trajectory inference field is beyond the scope of this paper. Instead, we restrict ourselves to discussing how existing velocity methods, as well as certain clustering algorithms, can be represented as special cases of the exact solution in Equation 23.

### 4.6 Clustering as a special case

First, consider the ergodic case of *f* on [0, *T*] with *T* → ∞, and suppose there are *K* cell types at equilibrium. These cell types are distinguished by gene-specific parameter vectors 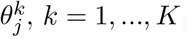. Assume only a single RNA species per gene exists, and all genes are independently expressed in a single cell type, without yet imposing a specific biological model. This yields the likelihood

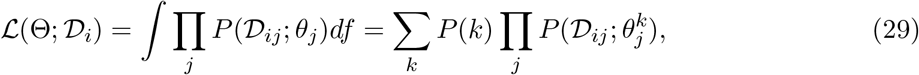

where *P* (𝒟_*ij*_; *θ*_*j*_) is the probability of observing the data 𝒟_*ij*_ and *P* (*k*) ∈ *θ*_*G*_ is the probability of cell *i* being in cell type *k*. The process time *t* is no longer necessary, because all cell types are stationary: the likelihoods are evaluated at the ergodic limit *t* = ∞. Equation 29 amounts to saying that the likelihood of a cell’s observation can be represented by using the law of total probability and conditioning on the cell type:

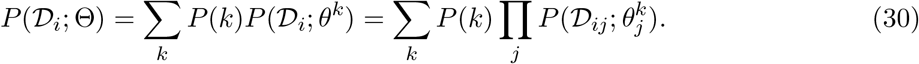

To optimize this likelihood, we need to specify *P* (𝒟_*ij*_; *θ*_*j*_), which is informed by the biophysics of transcription and mathematical tractability. The log-normal distribution is particularly common: if we treat log-counts ln 𝒟, the law of *P* (ln 𝒟_*j*_; *θ*_*j*_) is Gaussian. The lognormal distribution can emerge from several hypothesis: through a common, if *ad hoc* approximation to the gamma distribution which emerges from the mesoscopic limit of the CME [148], from the exact solution of a deterministic, macroscopic model with log-normally distributed transcriptional rates [149], and by mere assertion that the negative binomial distribution is similar to the lognormal distribution, without further discussion [147].

The lognormal approximation implies that each “cell type” is essentially a high-dimensional normal distribution in logarithmic state space. This induces a set of gene- and cluster-specific log-means 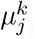, log-standard deviations 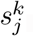, and a cluster assignment vector *σ*, such that *σ*_*i*_ ∈ {1, …, *K*}. The problem of characterizing cell types, i.e., fitting *P* (*k*), *μ*^*k*^, and *s*^*k*^, and providing an optimal point estimate of *σ*, under this model is equivalent to using the expectation-maximization algorithm to fit a Gaussian mixture model to the logarithmic data [150]. However, some caveats deserve mention. The model choice requires careful consideration. The log-normal heuristic is incoherent with the standard velocity model, which tends to a normal distribution with equal mean and variance in the continuous limit. Furthermore, although the Gaussian mixture model formulation can be justified as an approximation of a particular class of models, it is unlikely that this approximation holds generally.

For completeness, we note that the standard alternative to Gaussian mixture model clustering is community detection based on a graph constructed by defining a neighborhood criterion among cell vectors [36]. However, it has not yet been shown that such an approach can be afforded any well-defined probabilistic meaning. The literature contains numerous assertions that a meaningful Markovian transition probability matrix can be defined on observed cell states [1, 9, 10, 25, 138]. However, the constructed Markov chains have not been demonstrated to possess any particular relationship to an actual biological process.

### 4.7 The “deterministic” *velocyto* model as a special case

Strictly speaking, we only need to solve Equation 23 if we want to exploit useful properties of likelihood landscapes and estimators. However, if we are willing to forgo these advantages, we can use a moment-based estimate.

The linearity of the occupation measure can be used to compute summary statistics. For example, we can treat the RNA velocity model defined in Equation 1, with *μ*_*u*_(*t*) and *μ*_*s*_(*t*) giving the instantaneous process averages. The following relations hold at each instant *t* and over the entire trajectory:

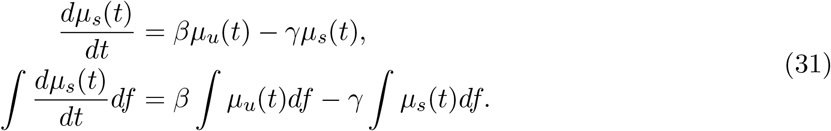

Each species’ mean occupation measure *μ*can be related to the instantaneous mean *μ*(*t*):

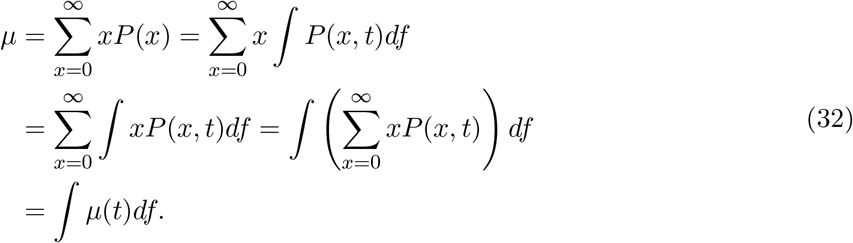

This implies the following identity:

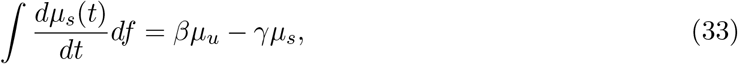

which holds regardless of the transcriptional dynamics encoded in *α*(*t*).

The identity formalizes the moment-based approximation to the biological parameters: if the left-hand side (the net velocity of the process) is sufficiently close to zero, the right-hand side gives an estimate for *γ/β*. Conversely, if this condition is violated, a naïve fit based on the moments (equivalently, a least-squares fit with zero intercept) will be biased by transient contributions, motivating the use of the extrema fitting procedure (as in Fig. 2 in SN2 of [1]).

We can investigate the behavior of the net velocity in a simple model system. Suppose *β* = 1, *df* = *T* ^−1^*dt*, and *α* is piecewise constant. We define *α*_*k*_, with *k* ∈ {1, …, *K*}, constant on each interval *I*_*k*_ ∈ [*τ*_*k*−1_, *τ*_*k*_]; the bounds are *τ*_0_ := 0 and *τ*_*K*_ := *T*. We define the length of an interval as Δ_*k*_ = *τ*_*k*_ − *τ*_*k*−1_. The following equations hold for every interval:

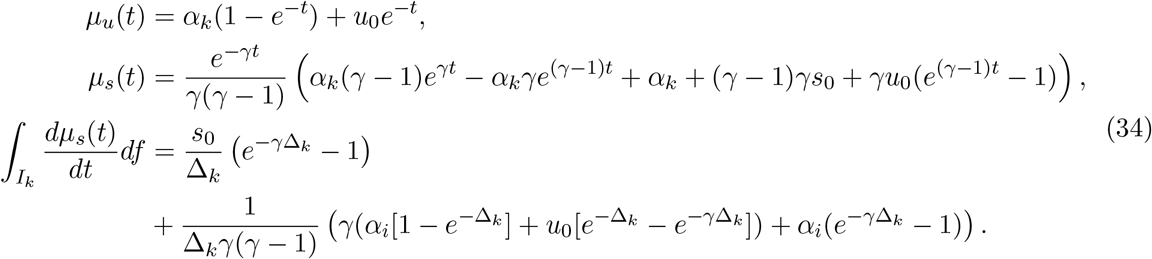

In each interval *I*_*k*_, the integral approaches zero as Δ_*k*_ grows. This result has a qualitative interpretation: as interval duration grows, the process settles into its ergodic equilibrium attractor, and that attractor provides an effective estimator of *γ*. On the other hand, if the interval is short-lived relative to mRNA lifetime, the integral is dominated by the initial condition, and the system is largely out of equilibrium.

In practice, this means that the degradation rate is identifiable through moments only if the lifetimes of the mRNA species are short relative to the interval lengths, i.e., the net velocity is low enough. On the other hand, if the lifetimes are too short, the transient regimes are sparse and steady states are approached rapidly, giving no information about dynamics (and reducing the problem to the formulation in Equation 29). The quantile fit procedure is simply a heuristic method to winnow the data for near-equilibrium populations under the informal prior that these populations are present in the data. Unfortunately, this approach is subject to the usual pitfall of moment-based methods: if the prior is wrong, there is no easy way to identify its failure.

We have omitted the discussion of normalization and imputation, as they are not amenable to analysis. However, their impact can be evaluated by generating data from the model and processing it with the *velocyto* workflow. The intuition outlined above concords with our simulations: in Figure 8e, we fit the simple model introduced in Section 4.3 using three different methods. In i., we use simple linear regression on the raw data; in ii. and iii. we apply a quantile fit to normalized and imputed data respectively. In spite of the distortions in the overall phase portrait (Figure 8c), the extrema are stable enough under imputation to generate fairly reliable estimates of *γ/β*, up to roughly an order of magnitude, whereas the ratio of averages is significantly less precise. This is consistent with the performance reported in Figure 5 (as in the *k* = 0 case, which uses linear regression on the entire dataset).

In light of the formalization, the fitting procedure is contingent rather than stable, necessitating careful study of limitations. Using the average of the full occupation measure is contingent on the net velocity being near zero (Δ_*k*_ sufficiently large for all *k*). Using the average of the extrema is contingent on those extrema having equilibrated (Δ_*k*_ sufficiently large for *k* with largest and smallest *α*_*k*_). Furthermore, it provides no information about the relative timescales *β*_*j*_ of different genes.

The “stochastic” model, which was introduced in *scVelo*, is practically identical, but exploits additional information from the second moments of the extremal observations. This approach inherits the same issues, such as the reliance on the existence of extrema, and omission of *β*_*j*_, and introduces new ones, such as the assumption of identical error terms for first and second moments and the inference of error covariance parameters. In principle, further investigation may characterize whether these issues improve or worsen moment-based inference. However, we suggest that these details are marginal compared to the more fundamental limitations, as well as the discrepancies observed in simulation (Figure 8d iii-iv).

### 4.8 The “dynamical” *scVelo* model as a special case

The modeling approach we have presented can be used to contextualize part of the “dynamical” algorithm proposed in *scVelo*. First, assume that cell type transition times *τ*_*jk*_ ∈ *θ*_*j*_, i.e., no global parameters shared by multiple genes exist (*θ*_*G*_ = ∅). This reduces Equation 23 to the following form:

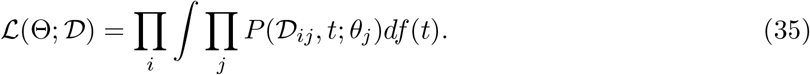

Omitting the uncertainty represented in the integral by assigning a time *t*_*i*_ to each cell we obtain:

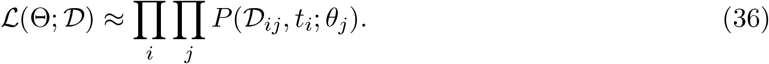

Since time assignments *t*_*i*_ are now deterministic, rather than probabilistic, likelihood landscapes become inaccessible. In principle, finding the maximum of Equation 36 can still provide a point estimate of Θ, although this may not be practical: now, *f* has to be inferred empirically by fitting *t*_*i*_. Applying the logarithm, we get

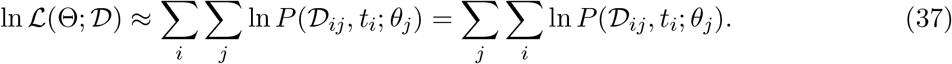

Using a Gaussian kernel centered on *μ*_*u*_ and *μ*_*s*_, with standard deviation *s*_*j*_ for both species, as an approximation to the likelihood, we can interpret *P* as a probability density function:

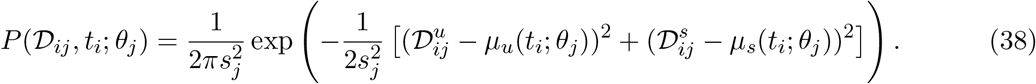

The question of what this kernel means, i.e., what biophysical phenomena it models, is subtler than it appears. On the one hand, there are certain regimes where stochastic spliced and unspliced counts are approximately distributed about the true averages *μ*_*u*_ and *μ*_*s*_ according to normal laws, such as in the high-concentration limit of certain jump processes explored by Van Kampen [142]. However, this limit yields time-dependent and concentration-dependent *s*_*j*_, incompatible with Equation 38 and the “stochastic” model described in the *scVelo* manuscript. Instead, this form presupposes *μ*_*u*_ and *μ*_*s*_ are the true deterministic amounts of mRNA, corrupted by Gaussian isotropic error without an explicitly named source. This model exemplifies the signal processing paradigm, which attempts to identify an underlying “signal” by regressing out incidental Gaussian “noise,” with various levels of biological justification [151–153].

For a particular gene, the log-likelihood of Equation 38 takes the following form:

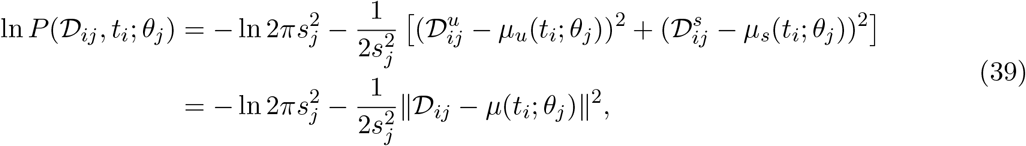

i.e., we can use a simple two-dimensional Euclidean norm to estimate the log-likelihood of the observation. For *M* independent observations of gene *j* with identical parameters *θ*_*j*_ but distinct times *t*_*i*_, we find that

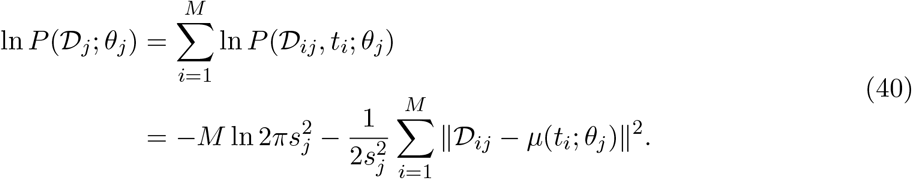

The optimum of this function is invariant under scaling, so we can work with the normalized negative log-likelihood:

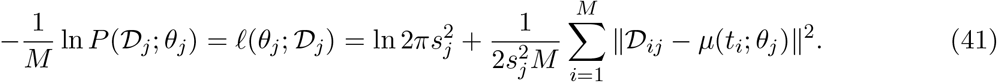

This contradicts Equations 7-8 of [3], which use a univariate, rather than bivariate, Gaussian error term. On the other hand, the actual implementation appears to use separate *s*_*j*_ for the two species, computed from the extremal points, and combine them in an *ad hoc* fashion.

In principle, solving Equation 37 with the likelihood indicated in 39, i.e., iteratively inferring *t*_*i*_ and *θ*_*j*_, yields a coherent maximum likelihood estimate of the system parameters. However, the method goes one step further, essentially taking Equation 37 and rewriting it in terms of the negative log-likelihood:

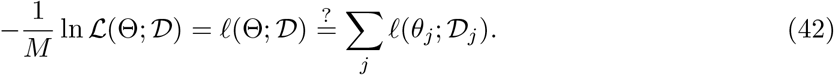

This approach posits that likelihood optimization over *N* genes be split into *N* independent problems, which can be parallelized. However, it is incorrect, as the times *t*_*i*_ are incoherent between different genes *j*, and the results are uninterpretable. This issue has been tacitly acknowledged, e.g., in a *post hoc* approach adopted to make times “agree” in *scVelo* [3] and a similar proposal by Li et al. [25], although without any rigorous justification.

The actual implementation does not use the raw data 𝒟_*ij*_. Rather, it uses a normalized and imputed version of the data. Again, the effect of these transformations is challenging to characterize analytically. However, the simulations shown in Figure 8c-d and Figure S5 suggest that there is no compelling reason to believe the imputed data reliably estimate the underlying averages *μ*_*u*_ and *μ*_*s*_ for a stochastic system out of equilibrium. Furthermore, the noise-corrupted deterministic model used in the likelihood computation is biologically implausible.

### 4.9 Prospects for inferential procedures

In Section 4.2, we have presented a framework for the description of transient stochastic systems. This framework is versatile enough to describe a range of problems in single-cell sequencing, from clustering to trajectory inference. The methods presented in previous RNA velocity publications are best understood as approximations to exact solutions under fairly strong, informal priors about the process biophysics. The *velocyto* algorithm uses a moment approximation, which assumes the system has effectively equilibrated. The *scVelo* algorithm uses a Gaussian-perturbed ODE model, which assumes the mRNA counts do not have any intrinsic noise, only isotropic measurement noise. The latter yields considerably more information from the data, but imposes considerably stronger assumptions, making the obtained information essentially uninterpretable.

However, our formalization immediately presents options for likelihood-based parameter estimation. We introduced an approximate method that does maintain cell identities and motivated it using the properties of order statistics. This method is challenging because it requires a fairly involved combinatorial optimization. However, it does lend itself to developing further approximations, and provides routes for falsifying hypotheses. We believe that such biophysical models, amenable to approximation and testing, are crucial for the future interpretation of dynamics predictions from sequencing data.

### 4.10 Embedding

To complement the internal controls discussed in Section 3.6, we performed a set of comparisons with data simulated with no unknown sources of noise. We embedded a simulation of the system introduced in Section 4.3 and illustrated in Figure 8 into a two-dimensional principal component space. The results are shown in Figure 9. Even the ground truth velocity arrows (Figure 9a) only retained a small amount of information after the transition from 100 dimensions to two. This experiment provides us with the answer to the second question in Section 3.2: even if we have “true” velocity directions, they only contain a limited amount of highly local information. As expected from the fair performance in Figure 8e iii, the inferred linear embedding (Figure 9b) was globally and locally (Figure 9f) faithful: the model precisely matches the assumptions of the parameter inference workflow. However, the estimates were rapidly distorted upon applying the nonlinear embedding procedure (Figure 9c), rotating many cell-specific directions and suggesting transitions from the green reverting population to the light blue perturbed population, whereas, the true trajectory is from light blue to green. The results of the Boolean procedure were slightly more faithful to the linear projection (Figure 9f) but otherwise qualitatively similar (Figure 9d). This is the method’s best-case performance.

Even in the PCA projection, the performance of the nonlinear velocity embedding leaves much to be desired: the procedure is biophysically uninterpretable, discards the vast majority of information, and risks failure when model assumptions are violated. For example, it can generate false positive velocity fields when the ground truth is completely static (Figure S6, as simulated using the procedure in Section 6.2); even directly inspecting the phase plots may be insufficient diagnose to this problem (e.g., compare Figure S7 to Extended Data Figs. 6c and 7c of [1] and Figs. 2c and 3g of [3]).

The nonlinear, non-deterministic embeddings ubiquitous in the analysis of scRNA-seq data degrade the performance further. In Figure 10, we embedded a system with three potential terminal states, generated by the simulation procedure described in Section 6.2. Cell projection into PCA appeared to conflate two of the branches; the nonlinear embeddings effaced causal relationships altogether. As before, the arrows were broadly coherent whether or not they included quantitative information. Finally, as described in Section 1.1, the embedding procedure has previously demonstrated catastrophic failure to capture known dynamics in biological datasets [5, 9, 10, 13, 22]. Therefore, although embeddings are qualitatively appealing, they are unstable, challenging to validate, and harbor intrinsic global- and local-scale pitfalls that arise even in simple scenarios.

**Figure 10:**
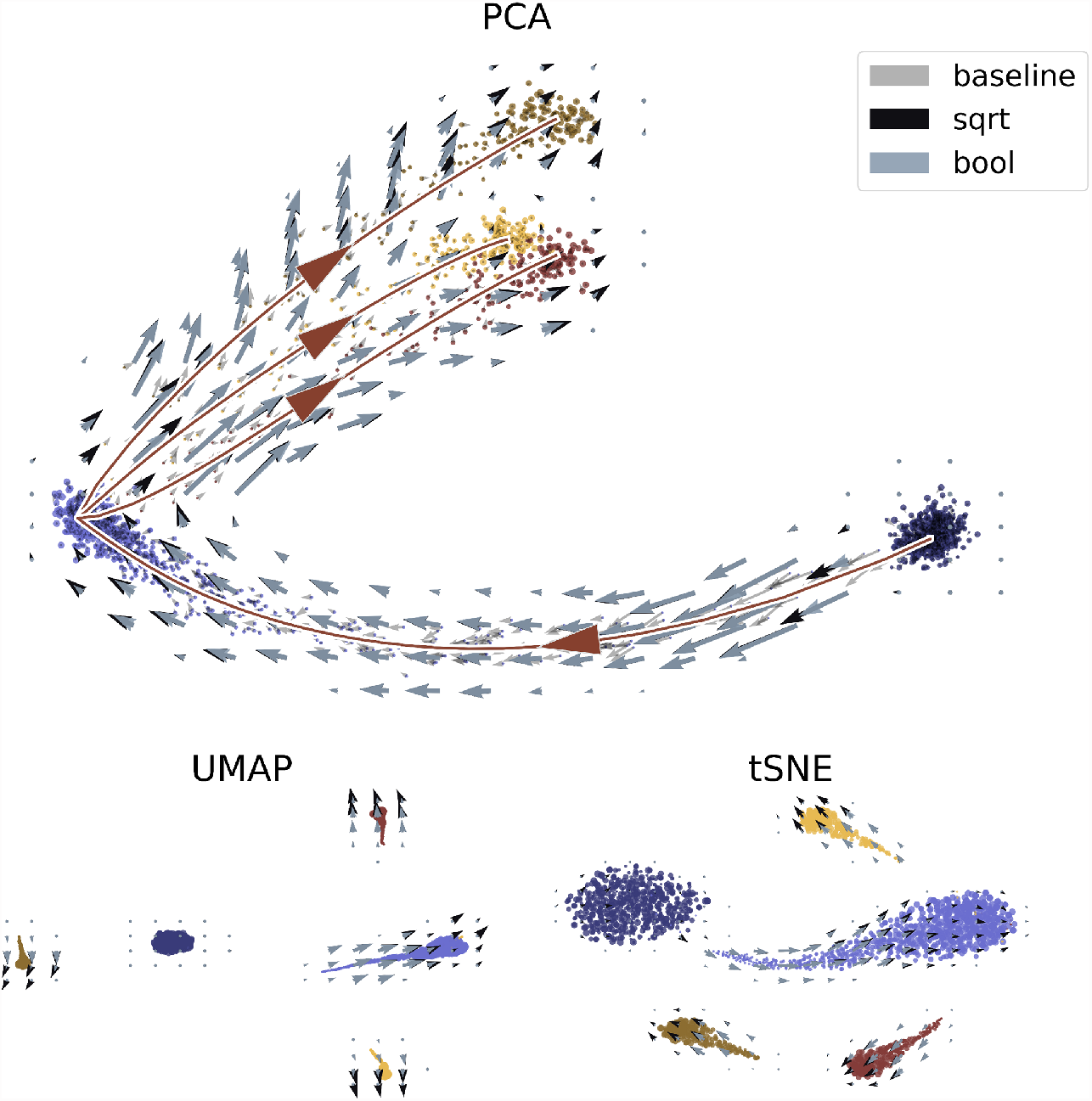
Performance of cell and velocity embeddings on simulated data, compared to ground truth principal curve. Top: PCA embedding with linear baseline and nonlinear aggregated velocity directions, as well as ground truth principal curve; trajectory directions displayed to guide the eye. Bottom: UMAP and *t*-SNE embeddings with nonlinear velocity projections.

Nevertheless, some human-interpretable visualization is desirable. In light of the frustrating dearth of theory and interpretability for the commonly used embedding procedures, we note that the stochastic formulation we have presented can be used to speculate about more rigorous and stable embedding methods that would use rather than discard quantitative information. Instead of individual cells, which inevitably exhibit noise, we suggest constructing and emphasizing the underlying graph governing parameter values (as in, e.g., Fig. 1F of [138]). Alternatively, since the current low-dimensional embeddings are used to support claims about presence of *a priori* human-interpretable features such as equilibrium cell types, limit cycles, and transient differentiation trajectories, it may be better to fit a hierarchical model consisting of those features and reporting the best-fit model. For example, if the goal is to cluster data, then it makes sense to fit Equation 29. On the other hand, if the goal is an elucidation of tree-like differentiation trajectories, it may be better to incrementally grow a trajectory mixture model until its complexity outweighs its likelihood per a statistical information criterion. Formally, this would correspond to optimizing the likelihood of samples from analogs of Equation 18. If a method has succeeded in inferring the underlying topology and dynamics, a meaningful and well-defined principal curve induced by the underlying mechanism, as shown in Figure 9e and the PCA in Figure 10, could be plotted.

## 5 Conclusion

### 5.1 Summary

The two main steps in RNA velocity, namely the model estimation and embedding, originate from different approaches to data analysis that can be at odds with each other. The count processing and inference steps, which comprise the model estimation procedure, serve to identify parameters for a transcription model under some fairly strong assumptions, such as constitutive production and approximately Gaussian noise. This procedure can be treated as an informal approximation of a method to solve a system implied by the simulation design in the original publication. However, as we have seen, this system abstracts away many aspects of the technical artifacts present in single-cell RNA sequencing, and the transcriptional dynamics that drive the molecular biology of cells. The embedding processes used, which are entirely *ad hoc*, discard nearly all of the quantitative information, and can occasionally fail. Particularly problematic is that the failures, when they occur, are difficult to identify. Moreover, failure may result from many problems, including overlaps in the embedding and erroneous clustering. Such problems may be mitigated or exacerbated by tuning hyperparameters. These challenges, contradictions, and the assumptions inherent in the many choices that are made for each of the steps, have not been previously characterized in full detail, and they add up to a mixed picture. On the one hand, at least in some simulated cases, the RNA velocity method does work, and the latent signal is strong enough to capture broad trends. On the other hand, catastrophic failure can lurk at any step of the velocity workflow, and there are no theorems to alert users to failure modes, or to diagnose or delimit the extent of failures. Instability and reliance on user-tuned hyperparameters are not grounds for abandoning the method; the same problems crop up with kernel density estimation, *k*-means clustering, histogram binning, time series smoothing, and many other analysis tasks. However, the tendency to compensate for lack of theorems with more *ad hoc* filters and more sophisticated modeling that requires optimization of neighborhood sizes, normalization procedures, and thresholds only exacerbates the problems.

The mathematical foundations of stochastic biophysics have been studied for several decades; they are well-understood, and amenable to generalizations and approximations. The chemical master equation allows for the elucidation of technical noise [128], and the quantitative exploration of transcriptional regulation [148] and splicing [78]. As discussed in Section 4.2, the same modeling framework can be used to describe general classes of differentiation processes. Rather than starting with heuristics and then seeking to unravel their meaning, in this approach one begins by motivating, defining, and solving a general system, and only subsequently deriving approximations and statistical summaries. These can range from simple moment expressions to low-dimensional principal curves as illustrated in Figure 10. Furthermore, with such an approach, one can leverage the machinery of Bayesian inference to directly fit full distributions, with the advantages of interpretability and statistical robustness. This highlights that the primary challenge in RNA velocity is not its extension via additional heuristics, but rather the development of tractable inference procedures.

### 5.2 Proposals

We conclude with a summary of the main steps of an RNA velocity workflow, along with some insights and proposals that emerged from our work:

#### Pre-processing

As demonstrated in Section 3.1, the specific choice of processing software does produce discrepancies in results, although the small, and largely arbitrary, variations in assignment rules do not provide compelling reasons to select one method over another, pending the development of more detailed splicing network definitions. There are, however, substantial processing time differences [28,29] that can affect reproducibility of results, leading us to prefer fast pseudoalignment methods.

#### Filtering

This step is necessary for tractability, since full spliced and unspliced matrices can be challenging to analyze. While caution must be applied in selecting thresholding criteria, we find no reason to deviate from the standards typically applied in RNA velocity analyses.

#### Model definition

RNA velocity methods have been inspired by stochastic models of transcription. However there has not been a strong link between the models and the implemented methods, which are based on loose analogies and heuristics. We believe that explicit construction and discussion of biophysical models is imperative when developing RNA velocity methods, so that results can be meaningful and interpretable. In particular, we caution against the class of continuous and constitutive models implemented in velocity packages thus far; as discussed above, bursty models are tractable [78] and substantially more plausible according to live-cell data.

#### Normalization and imputation

The normalization and averaging of data to produce continuous curves is intended to remove cell size effects and to denoise the data. We found several problems with this approach. Firstly, this assertion is not motivated by theory, and our theoretical concerns in Section S1.1 suggest that model-agnostic “correction” is inappropriate. Secondly, as discussed in Section 4.3 and illustrated in Figure 8, the imputed data do not accurately recapitulate the supposed ground truth even in the simplest case. Finally, imputation prevents the most natural interpretation of counts as discrete random variables. Based on Kim et al. [108] and [128], we advise against normalization; it is more meaningful and accurate to apply parameterized models of extrinsic noise and gene–gene coupling. The interpretability afforded by discrete models outweighs the potential benefits of *ad hoc* normalization. Furthermore, we strongly recommend against imputation more generally: studies such as [65, 108, 154] have revealed distortions, and the approach possesses fundamental instabilities (as in Figure 5).

#### Inference

From a probabilistic perspective, current inference procedures are problematic. Instead of currently implemented procedures, it is more appropriate to build and solve mechanistic, fully stochastic models that allow for fitting copy numbers. This can be computationally facilitated by a data selection process coherent with the “marker gene” paradigm: if a gene does not need to be fit to a transient model, one should not try to fit one. Thus, we recommend fitting ergodic distributions to genes that are not meaningfully modulated across the dataset, ergodic mixture distributions to (fewer) genes that vary across disjoint cell types, and occupation measures to (even fewer) genes that exhibit transient behaviors. Although joint inference is relatively challenging, we believe that a formulation that can exploit existing combinatorial optimization frameworks may be a productive avenue for exploration.

#### Embedding

RNA velocity embedding procedures inherit problems accrued with the steps discussed above. However, even in an idealized situation where an interpretable and well-fit model is used, current embedding practices are counter-productive for interpreting the data, as discussed with reference to controls in Sections 3.6 and 4.10. Despite the implication of causal relationships between cells encoded by cell–cell transition probabilities, embedding procedures are currently *ad hoc*. Moreover, it has been shown that current methods distort local neighborhoods and the global topology in an unpredictable manner [120,122]. Directed graphs over the cells are attractive, but do not have a coherent interpretation relative to the underlying biophysics. Instead of such methods, we recommend directly working with the latent process governing the transcriptional variation. Nevertheless, two-dimensional visuals may be useful for summarizing the raw data; using simulations, we demonstrate an interpretable method for embedding a true principal curve in deterministic principal component space in Section 4.10.

## 6 Methods and Data

### 6.1 Pre-processing concordance

There is no well-defined “ground truth” for mRNA counts in arbitrary datasets. However, to obtain a qualitative understanding of potential pitfalls, we performed controlled experiments to analyze the discrepancies between outputs produced by popular software implementations.

The concordance analysis was heavily inspired by the benchmarking of Soneson et al. [28]; however, our goals and scope differ. First, we sought to analyze the reproducibility of the findings across several datasets; the original analysis only treated a single dataset generated using the 10x v2 chemistry. To this end, we analyzed ten datasets that used the 10x Genomics v2 and v3 protocols. Second, we sought to focus on the processing workflows most relevant to casual use. The original analysis examined thirteen quantification workflows, whereas we examined three: *velocyto, kallisto*|*bustools*, and *salmon*. These were run with default settings. We have made available all the scripts and *loom* files generated by the workflows (Section 6.9).

We obtained ten datasets generated with the 10x Genomics scRNA-seq platform. Two were released as part of a study by Desai et al. and used v2 chemistry [155]. Eight were released by 10x Genomics and used v3 chemistry. The dataset metadata are outlined in Section 6.9.

To implement the *velocyto* workflow, we ran *CellRanger* on the datasets using human and mouse reference genomes, pre-built by 10x Genomics (*GRCh38* and *mm10* 2020-A). We then processed the aligned outputs using the `run10x` command provided in *velocyto*.

To implement the *kallisto*|*bustools* workflow, we ran the `ref` command on the pre-built genomes to build references, using the standard `--workflow lamanno` option. We then processed the raw data with the `count` command, passing in the generated reference and using the `--workflow lamanno` option.

To implement the *salmon alevin-fry* workflow, we ran the *alevin-fry* velocity workflow documented at https://combine-lab.github.io/alevin-fry-tutorials/2021/alevin-fry-velocity/, from the initial reference construction to the final *anndata* output. This output was converted directly to loom files for the comparative analysis. We used the same pre-built 10x Genomics reference genomes (*GRCh38* and *mm10* 2020-A) as above.

### 6.2 Simulation

#### Transient constitutive model: perturbation and reversion

To generate Figure 8, we simulated data from the constitutive transcription model with the “cell type” structure *ABA*. In this model all cells start out in state *A* at *t* = 0, switch to state *B* at *t* = *τ*_1_, and revert back to state *A* at *t* = *τ*_2_. We generated 2000 cells and 100 genes. As shown in Figure 8a and formalized in Equation 21, we defined three time periods corresponding to each cell type. The simulation time horizon was set to *T* = 10, with synchronized transition times *τ*_1_ = 3 and *τ*_2_ = 7. The gene-specific transcription rates *α*_1_ and *α*_2_ were generated from a lognormal distribution with log-mean 0 and log-standard deviation 1. The gene-specific splicing rates *β* were generated from a lognormal distribution with log-mean 1 and log-standard deviation 0.5. The gene-specific degradation rates *γ* were generated from a lognormal distribution with log-mean 0.5 and log-standard deviation 0.25, to reflect the intuition that splicing is somewhat faster than degradation. Sampling times were generated from a continuous uniform random variable on the interval [0, *T*].

The solutions in Figure 8b were computed from an approximation to the generating function. We did not account for the initial condition, which suffices because the transcription rate on (0, *τ*_1_) is low. The true values of *μ*_*u*_ and *μ*_*s*_ were computed from the solutions to the governing ordinary differential equation. The true values of 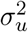 and 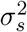 were set to *μ*_*u*_ and *μ*_*s*_, respectively, as the mean of a Poisson distribution is identical to its variance. To generate Figure 9, we simulated data from the same model, with 2000 cells and 100 genes.

#### Transient constitutive model: multipotent differentiation

To generate Figure 10, we simulated data from the constitutive transcription model with the “cell type” structure *AB*(*C/D/E*). In this model all cells start out in state *A* at *t* = 0, switch to state *B* at *t* = *τ*_1_, and then transition to one of the terminal states *C, D*, or *E* at *t* = *τ*_2_. We generated 2000 cells for 100 genes. The gene parameter and observation time distributions, as well as switching times, were identical to those reported in Section 6.2. Each cell fate was assigned randomly, with equal probabilities of 1*/*3. We used an identical procedure for Figure S1.

#### Steady-state bursty model

To generate Figures S6 and S7, we generated synthetic data assuming that RNA transcription is at an equilibrium, but has heterogeneity due to different cell types. The bursty transcription model (Equation 19) was implemented using the PGF schema in Equation 20, using the approach outlined in its derivation [104]. We specified parameters for 100 genes, with cell-independent burst sizes *b* and splicing rates *β*. Burst sizes *b* were generated from a lognormal distribution with log-mean 0.3 and log-standard deviation 0.8, clipped to stay in the range [0.05, 25]. The splicing rates *β* were set to 1 with no loss of generality. We simulated 10 cell types distinguished by average burst frequencies *α* and degradation rates *γ*, with 300 cells per cell type.

Average gene-specific log-degradation rates ⟨*γ*⟩ were generated from a normal distribution with mean −0.3 and log-standard deviation 0.3, to reflect the intuition that splicing is somewhat faster than degradation [1]. Gene- and cell type-specific degradation rates *γ* were generated from a lognormal distribution with log-mean ⟨*γ*⟩ and log-standard deviation 0.1, clipped to stay in the range [0.08, 4], to reflect the intuition that extrinsic noise in degradation rates is low relative to that in transcription rates.

Burst frequencies were generated from a lognormal distribution with log-mean −1 and log-standard deviation 0.5, clipped to stay in the range [0.005, 1], to encode the intuition that transcriptional activity is relatively rare. Analytical means *μ*_*u*_, *μ*_*s*_ and standard deviations *σ*_*u*_, *σ*_*s*_ were computed for spliced and unspliced distributions. Histograms were generated up to *μ*+ 5*σ* in each direction, clipped to be no lower than 10. To keep molecule counts realistically low and the histograms tractable, we rejected and regenerated parameter sets that produced (*μ*_*u*_ + 5*σ*_*u*_) × (*μ*_*s*_ + 5*σ*_*s*_) > 1.5 × 10^4^. To generate observations, we sampled directly from the histograms.

The phase plots displayed in Figure S7 were manually selected after sorting for simulated genes with the highest Σ_*i*_ Δ*s*_*i*_ and −Σ_*i*_ Δ*s*_*i*_.

### 6.3 Filtering

In Figures 5-7, we analyzed the forebrain dataset. To pre-process it, we implemented a procedure largely identical to that used to generate Figure 4a of the original publication [1]. We performed several rounds of filtering:

For the forebrain dataset, we used the following sequence of thresholds:

1. Discarding cells in the 0.5th percentile of total unspliced counts.
2. Discarding genes with fewer than 40 spliced counts, or expressed in fewer than 30 cells.
3. Selecting the top 2000 genes by coefficient of variation vs. mean, as implemented in the *velocyto* function `score_cv_vs_mean`, with maximum expression average of 35 (based on spliced counts).

In all other figures, we analyzed simulated data and omitted filtering, as all genes *a priori* had the correct dynamics.

### 6.4 Normalization

After importing forebrain data, we normalized and log-transformed spliced and unspliced counts using the default schema implemented in the *velocyto* function `normalize`:

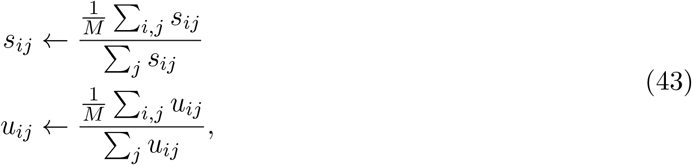

i.e., the “cell sizes” or total counts of spliced and unspliced molecules were separately normalized so each cell’s total was set to the mean over the dataset.

For the simulated data, we did not use normalization. This approach was inconsistent with previous simulated benchmarks [1], but we had three reasons for omitting it. First, as discussed in Section 4.3, normalization purports to “regress out” systematic technical and biological effects. We did not include these phenomena in the model. Second, the ground truth principal curves we constructed for the analyses in Section 4 (e.g., in Figure 9e) relied on evaluating the true gene-specific *μ*_*s*_(*t*) on a grid over [0, *T*], then log-transforming and projecting them to the two-dimensional principal component space. This was straightforward to do when the PC space was computed from raw counts, but more challenging otherwise. Finally, our omission made the velocity embedding procedure coherent: with the PC projection based on raw counts, we used the same underlying space to imputed counts and extrapolate velocities.

### 6.5 Embedding construction

For the forebrain dataset, we used size- and log-normalized spliced counts to construct the principal component projection. The UMAP and *t*-SNE embeddings were calculated from the top 25 principal components (as illustrated in Figure 6). For the simulated data, we used the log-normalized spliced counts to compute the same embeddings.

### 6.6 Imputation

Prior to fitting the models, data were smoothed by pooling across 50 nearest neighbors in the 25-dimensional principal component space constructed in Section 6.5, as quantified by Euclidean distance. To implement this step, we used the default parameters of the *velocyto* function `knn_imputation`. The choice of neighborhood space was arbitrary, and we imposed it for consistency. The original report used an adaptive principal component space based on the fraction of explained variance, whereas *scVelo* uses a default of 30 principal components. We observed no substantial difference in results between the adaptive and fixed schema. For the forebrain dataset, we pooled the normalized counts. In case of the simulations, we pooled the raw counts. In Figures 8 and S5, we deviated from this procedure to investigate the impact and suitability of normalizing simulated data. The figures demonstrate the respective effects of pooling the normalized and raw counts.

### 6.7 Inference and extrapolation

By default, the parameter *γ/β* was fit to the extrema of the imputed dataset: imputed unspliced counts were regressed as a linear function of imputed spliced counts with an offset. The extrema selection procedure used the defaults implemented in the *velocyto* function `fit_gammas`.

In Figure 8e, we deviated from this procedure to investigate the suitability and stability of inference from pooled data. In Figure 8e i, we performed linear regression on the raw counts of the entire dataset, whereas in Figure 8e ii, we performed linear regression on the quantiles of the normalized dataset. Finally, in the “Raw” or *k* = 0 cases illustrated in Figure 5, we performed linear regression on the raw counts of the entire dataset to contrast with regression on the extrema.

The standard inference procedure produced two parameters per gene *j*: the slope, a putative estimate of *γ/β*, and the intercept, which we denote as *q*. To compute the velocity of gene *j* in cell *i* for the nonlinear velocity embedding, we used the following formula:

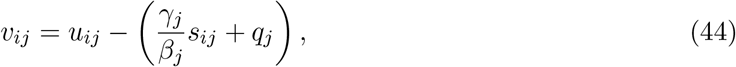

where *u* and *s* denote imputed quantities. To extrapolate the velocity and predict the spliced abundance after a time interval Δ*t*, we calculated Δ*s*_*ij*_ = *v*_*ij*_Δ*t*. This time interval was set to 1, for consistency with the *velocyto* implementation. The extrapolated value *s*_*ij*_ + Δ*s*_*ij*_ does not appear to be used in *velocyto*, as Δ*s*_*ij*_ contains the directional information used in the nonlinear embedding.

The schema described above is consistent with that of *velocyto*, but cannot be used to compute linear velocity embeddings (as given in Equation 3). The initial condition (*u*_*ij*_, *s*_*ij*_) used for extrapolation must be in the space used to build the PCA representation. Therefore, for linear velocity embeddings, we used Equation 44 in the corresponding space: normalized counts for the forebrain dataset and raw counts for simulated data.

However, if Δ*s*_*ij*_ < 0, the naïve extrapolation *s*_*ij*_ + Δ*s*_*ij*_ was not guaranteed to give a non-negative value that could be log- and PCA-transformed. In principle, we could have used an arbitrary Δ*t*, and clip any *s*_*ij*_ + Δ*s*_*ij*_ < 0 to zero. However, this approach, though closer in spirit to *velocyto*, would have risked extrapolating beyond the physical regime and could have introduced biases. Instead, we chose an extrapolation time Δ*t*^*^ guaranteed to stay in the physical regime:

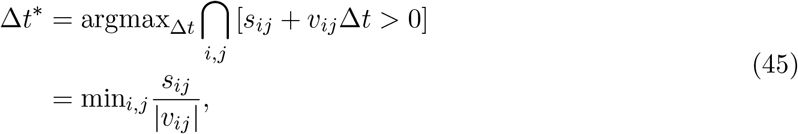

where we filtered for *v*_*ij*_ < −10^−6^. Finally, we set Δ*s*_*ij*_ to zero whenever *s*_*ij*_ < 10^−6^ and *v*_*ij*_ < −10^−6^, to avoid extrapolation into the negative regime: this fairly rare case occurs due to nonzero *q*_*j*_.

### 6.8 Velocity embedding

For the nonlinear velocity embeddings, we used 150 embedding neighbors and the square-root transformation by default. We deviated from this procedure in Figure S4 to investigate the impact of transformation and neighborhood choices. We used the default hyperparameter *σ* = 0.05 to calculate the softmax over directions to embedding neighbors (as described on pp. 7-8 in SN1 of [1]).

To implement the embeddings, we called the *velocyto* functions `estimate_transition_prob` and `calculate_embedding_shift`, which automatically correct for cell density. We did not use the neighborhood downsampling, randomization, or expression scaling options for the figures in this report; we observed no substantial difference in results between these schema and our standard procedure. The “high-dimensional space,” used to evaluate the displacements *s*_*q*_ − *s*_*i*_ in Equation 4, was the matrix of imputed spliced counts. The extrapolations Δ*s*_*i*_ were obtained from the procedure in Section 6.7.

For the linear velocity embeddings, we log- and PCA-transformed the matrix *s*_*ij*_ + *v*_*ij*_Δ*t*^*^, with the timescale obtained by the procedure in Equation 45.

The “Boolean” schema for velocity embedding is qualitatively similar to the schema proposed in the original publication; we previously proposed it to bypass the unit inconsistency between different genes’ *β*_*j*_ (and thus vectors *v*_*j*_) in the context of the *protaccel* package [4]. Instead of computing a correlation coefficient, we simply calculated the fraction of concordant signs between the velocity and the displacements to neighbors. In the parlance of Equation 4:

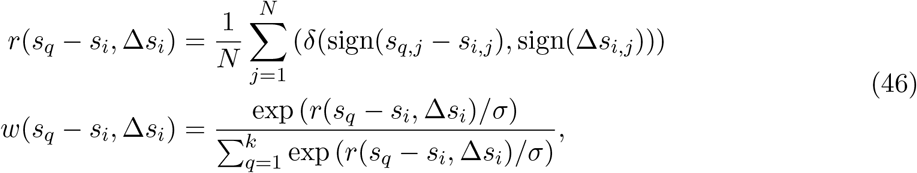

where *δ* is the Kronecker delta operating on inputs in {−1, 0, 1}.

We plotted cell-specific arrows for the linear baseline and aggregated the nonlinear velocity arrows using a 20 × 20 grid. We used this convention to distinguish the embedding methods, which are conceptually and quantitatively different, in plots that showed several velocity fields at once (e.g., the PCA plot in Figure 7). The grid directions were computed using the *velocyto* function `calculate_grid_arrows`, which applies a Gaussian kernel to average over the cell-specific embedded velocities nearest the grid point. We used the default parameters for the kernel, with 100 neighbors and a smoothing parameter of 0.5. We aggregated linear velocity projections in Figure 9a-b. As the arrow scale did not appear to have a quantitative interpretation, we set it manually to match the plot proportions.

### 6.9 Data availability

The datasets analyzed for Section 3.1, as outlined in Section 6.1, are listed in Table 1. The datasets released by Desai et al. were collated from the Sequence Read Archive (runs SRR14713295 for `dmso` and SRR14713295 for `idu`) [155]. The datasets released by 10x Genomics were obtained from https://support.10xgenomics.com/single-cell-gene-expression/datasets. The processed human forebrain dataset generated by La Manno et al. [1] was obtained from http://pklab.med.harvard.edu/velocyto/hgForebrainGlut/hgForebrainGlut.loom, as used in the *velocyto* documentation.

**Table 1:**
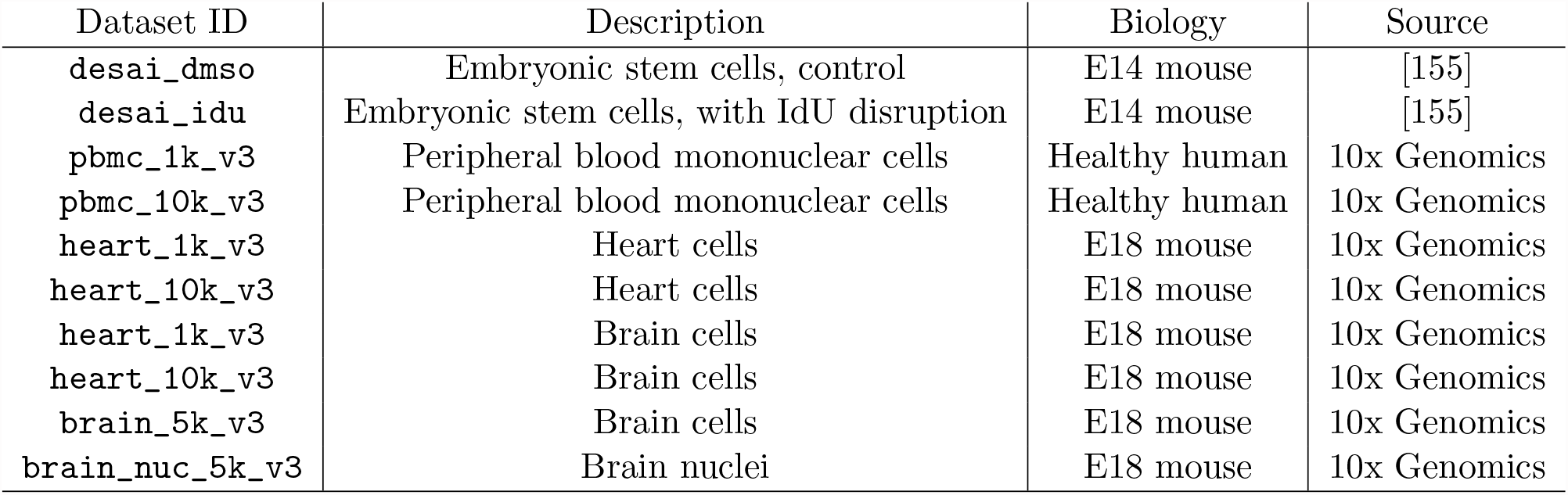
The datasets used to compare performance of molecule quantification software.

The processed *loom* files generated by the three workflows are available at the CaltechData repository, at https://data.caltech.edu/records/20030.

All Python scripts and notebooks necessary to reproduce the results of this study are available at https://github.com/pachterlab/GFCP_2022.

## 7 Author Contributions

Conceived of the project: G.G. and L.P.

Wrote scripts/notebooks for pre-processing, simulation, and analysis: G.G., M.F., and T.C.

Analyzed and interpreted the data: G.G., M.F., T.C., and L.P.

Wrote and edited the manuscript: G.G., M.F., T.C. and L.P.

## 8 Acknowledgments

G.G., M.F., T.C., and L.P. were partially funded by NIH U19MH114830. The DNA and RNA illustrations used in Figure 3 are derived from the DNA Twemoji by Twitter, Inc., used under CC-BY 4.0. The palette used in Figures 6, 8, 10, S3, and S5 is derived from dutchmasters by EdwinTh. G.G. thanks Dr. John J. Vastola for fruitful discussions about landscape representations of biophysical systems.

## S1 Supplementary Derivations

### S1.1 Model-agnostic count correction is impossible

In the body of this report, we discuss problems with the heavy-handed data treatment in the velocity workflows. In Section 3.4, we point out that it is inappropriate to fit an ODE model to normalized, pooled data: doing so amounts to presupposing a result that should be proven using careful theoretical analysis. In Section 4.3, we look for at least informal signs of this result in simulated data, and find nothing of the sort: imputation makes plausible-looking trajectories that nevertheless have very little in common with the ground truth. In the current supplement, we point out a much more general, yet elementary result that implies that count “correction” through imputation can be arbitrarily wrong.

Suppose we have a data point 𝒟_*ij*_ and the corresponding true mRNA abundance *n*_*ij*_ for a particular molecular species, cell *i*, and gene *j*. Sequencing is not perfect: the data point 𝒟_*ij*_ is generated from *n*_*ij*_ according to a non-deterministic schema, with an unknown probability law *P* (𝒟_*ij*_|*n*_*ij*_).

Two problems emerge. First, a point estimate of *n*_*ij*_ based on observed 𝒟_*ij*_ is necessarily incomplete: the sequencing process induces an entire distribution of possible *n*_*ij*_. This distribution is given by Bayes’ formula:

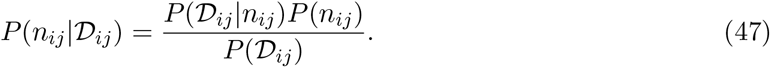

Assigning a single value is questionable, and downplays the effects of uncertainty. This remains a problem even if a theoretically optimal choice is taken, such as the point estimate 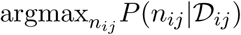.

Second, Equation 47 depends on *P* (*n*_*ij*_), the actual ground truth distribution. This distribution is unknown and needs to be identified and fit based on the data. Therefore, an imputation procedure that assigns a point estimate without considering the underlying distribution is *a priori* distortive.

In other words, this Bayesian argument illustrates that meaningful count correction is impossible without identifying and fitting the data-generating model, which encodes biological effects in *P* (*n*_*ij*_) and technical effects in *P* (*n*_*ij*_|𝒟_*ij*_). Count correction is strictly less powerful than parameter estimation for the biological and technical models, because count correction requires those parameters, whereas knowledge of the parameters immediately implies the entire distribution of the biological and observed variables.

## S2 Supplementary Figures

**Figure S1:**
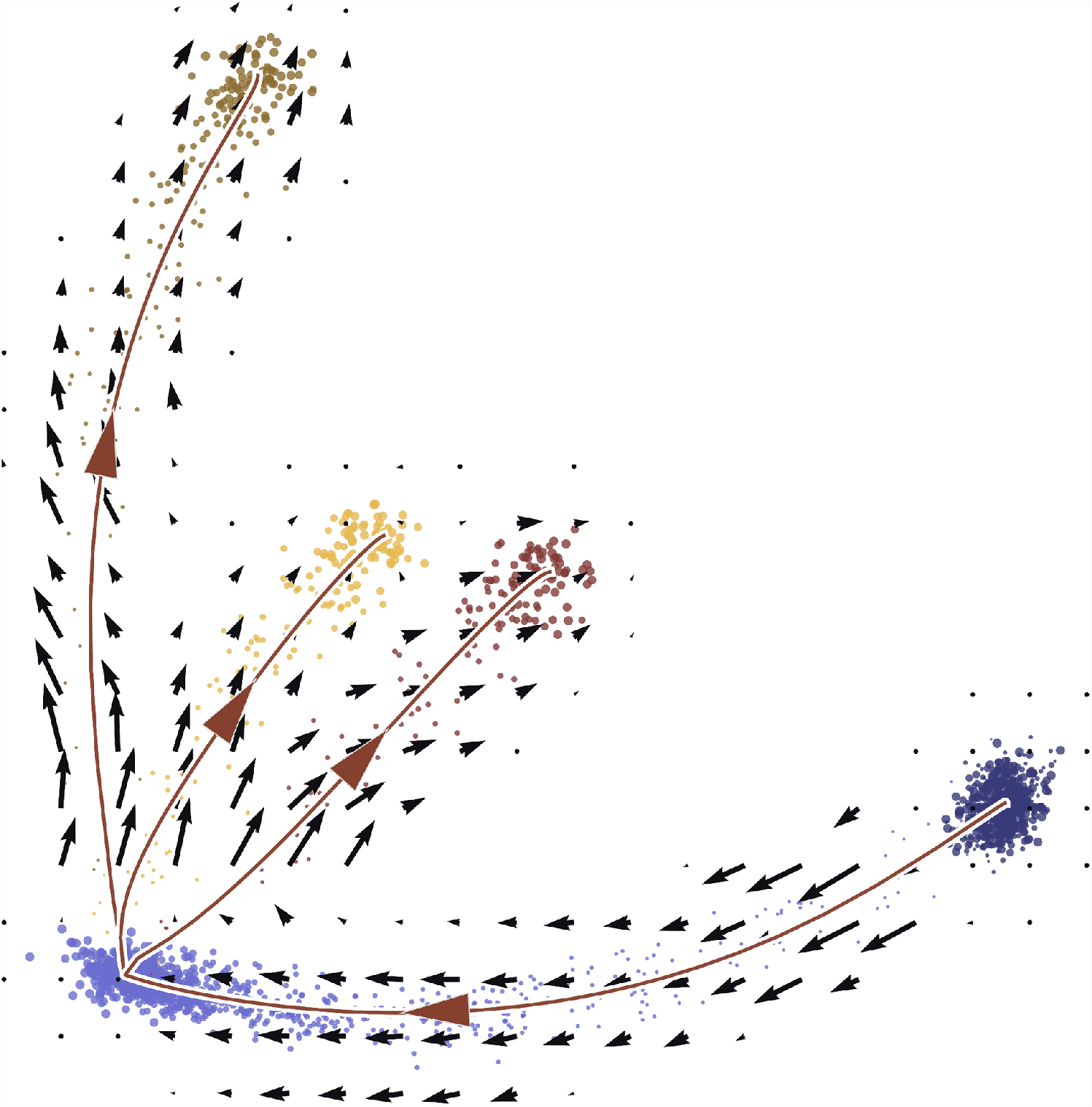
RNA velocity, as implemented in *velocyto*, can recapitulate the differentiation trajectories latent in data simulated from a tripotent trajectory. Trajectory directions are displayed to guide the eye (dark blue: cells in the source state; light blue: cells in the intermediate state; yellow, brown, and dark red: cells in one of the terminal states; black arrows: velocity embeddings produced using the standard *velocyto* workflow; lines with arrows: ground truth principal curves).

**Figure S2:**
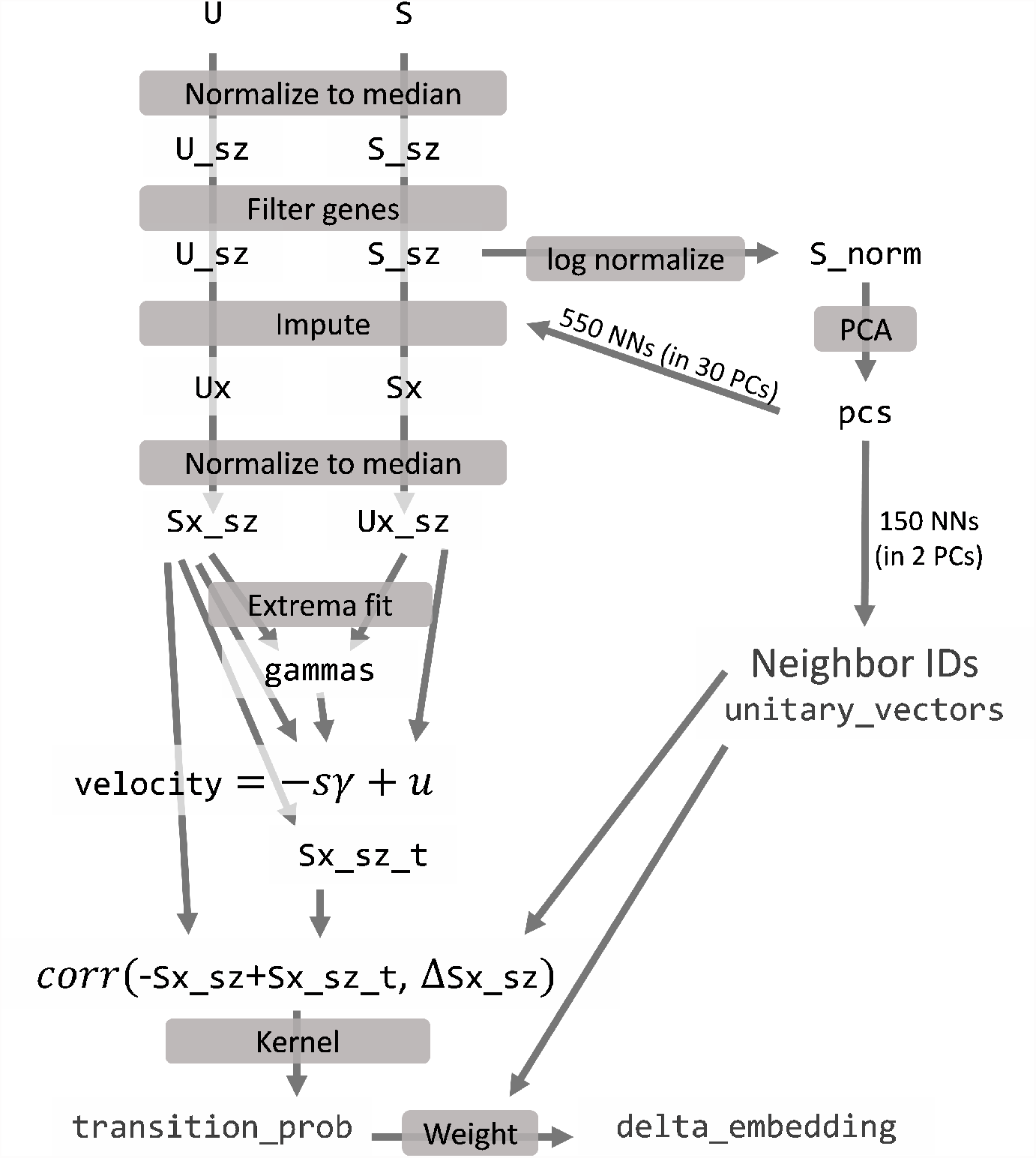
A summary of the data manipulations performed in a single run of the *velocyto* workflow.

**Figure S3:**
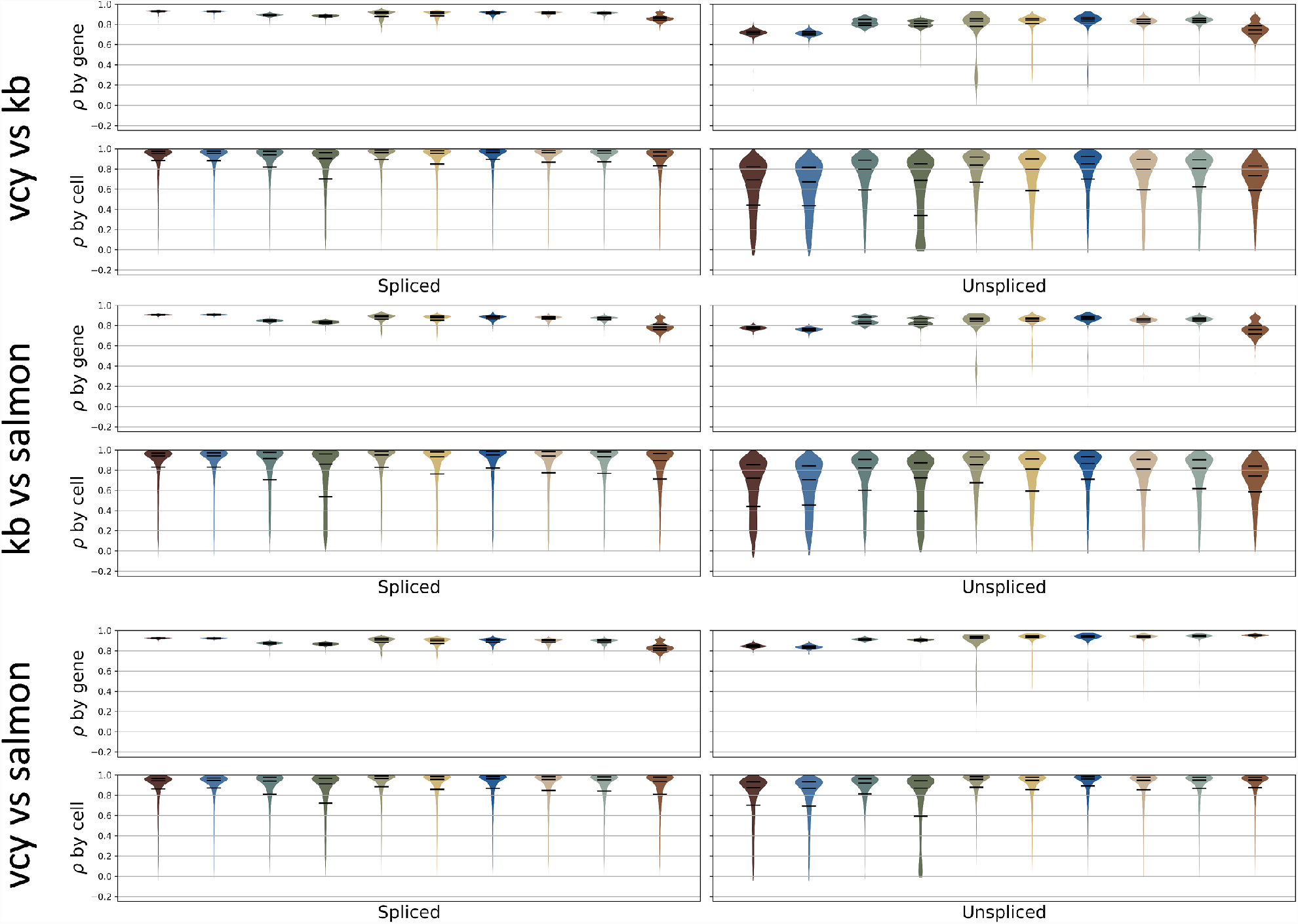
Distribution of Spearman correlation coefficients between outputs of *velocyto* (vcy), *kallisto*|*bustools* (kb), and *salmon*, as obtained from the 10 datasets enumerated in Section 6.9. Calculation “by cell” considers each gene in turn and computes correlation over all cells. Calculation “by gene” considers each cell in turn and computes correlation over all genes. Cells that only occur in a single software output and genes observed in fewer than four cells are omitted from analysis (Color: dataset; *ρ*: Spearman correlation value; violin plot: kernel density estimate of correlation distribution; horizontal lines: 25th percentile, median, and 75th percentile of correlation distribution).

**Figure S4:**
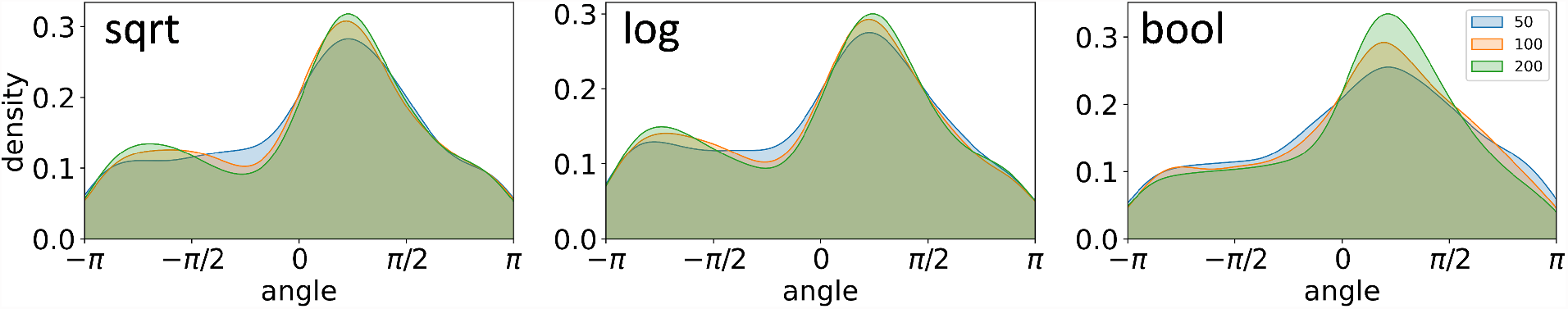
Nonlinear transformations and modulation of neighborhood sizes introduce distortions in the arrow directions with respect to the simplest PCA projection (subplots: different transformation procedures applied before kernel density estimation; histograms: distribution of cell-specific angle deviations under different pooling neighborhood sizes *k*).

**Figure S5:**
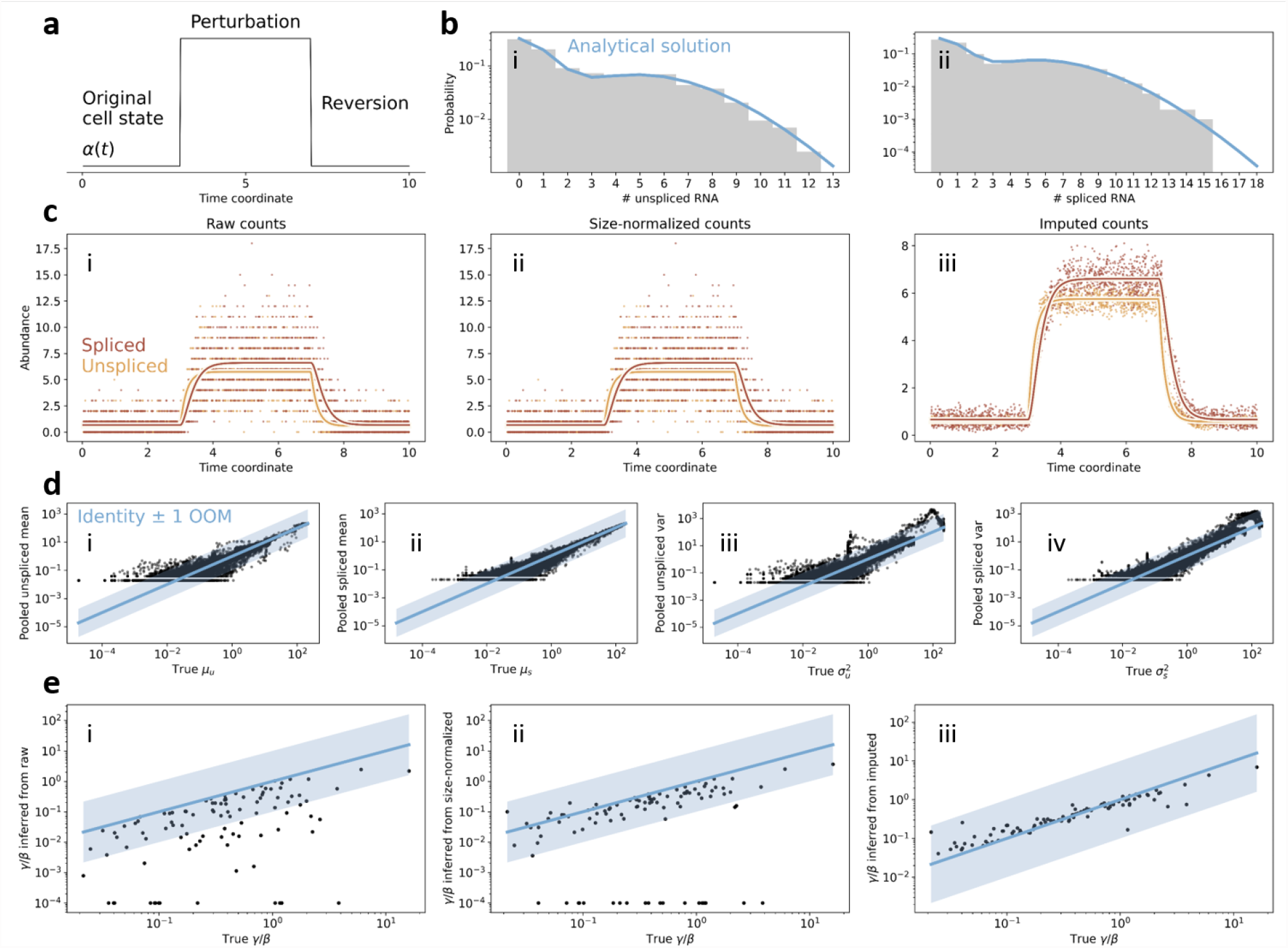
The RNA velocity count processing and inference workflow, applied to data generated by stochastic simulation, as in Figure 8 but without normalization with respect to the total number of molecules in the simulated cell.

**Figure S6:**
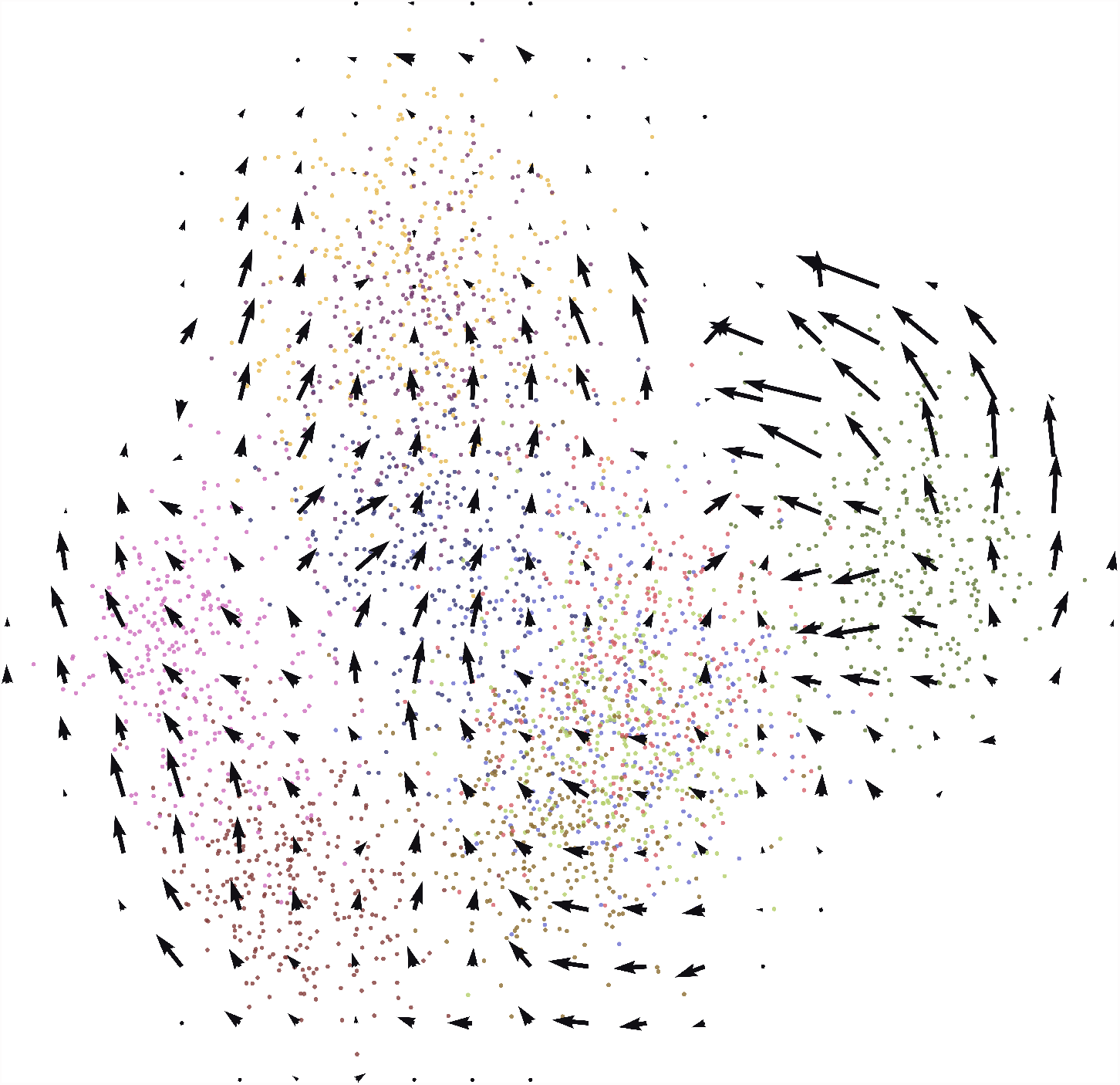
Erroneous velocity arrows resulting from a set of ten disjoint cell types with simulated bursty transcription (colors: ground truth cell types; arrows: velocity directions from default nonlinear embedding procedure).

**Figure S7:**
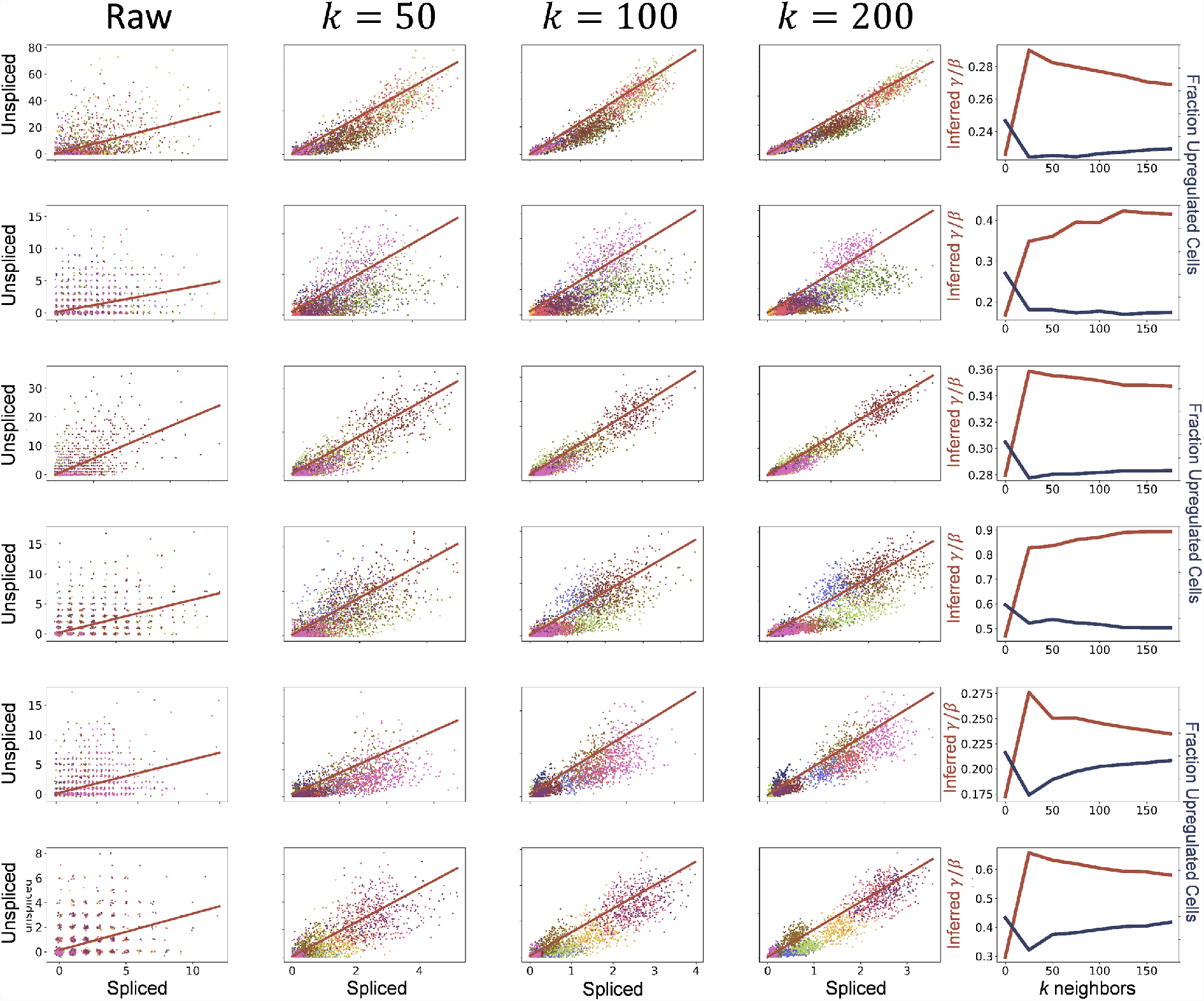
Phase plots resulting from applying the standard *velocyto* count processing workflow to data consisting of a set of disjoint cell types with bursty transcriptional dynamics (as in Figure S6).

